# A Structural Connectivity Atlas of Limbic Brainstem Nuclei

**DOI:** 10.1101/2022.08.10.503282

**Authors:** Simon Levinson, Michelle Miller, Ahmed Iftekhar, Monica Justo, Daniel Arriola, Wenxin Wei, Saman Hazany, Josue Avecillas-Chasin, Taylor Kuhn, Andreas Horn, Ausaf Bari

## Abstract

**Background:** Understanding the structural connectivity of key brainstem nuclei with limbic cortical regions is essential to the development of therapeutic neuromodulation for depression, chronic pain, addiction, anxiety and movement disorders. Several brainstem nuclei have been identified as the primary central nervous system (CNS) source of important monoaminergic ascending fibers including the noradrenergic locus coeruleus, serotonergic dorsal raphe nucleus, and dopaminergic ventral tegmental area. However, due to practical challenges to their study, there is limited data regarding their *in vivo* anatomic connectivity in humans.

**Objective:** To evaluate the structural connectivity of the following brainstem nuclei with limbic cortical areas: locus coeruleus, ventral tegmental area, periaqueductal grey, dorsal raphe nucleus, and nucleus tractus solitarius. Additionally, to develop a group average atlas of these limbic brainstem structures to facilitate future analyses.

**Methods:** Each nucleus was manually masked from 197 Human Connectome Project (HCP) structural MRI images using FSL software. Probabilistic tractography was performed using FSL’s FMRIB Diffusion Toolbox. Connectivity with limbic cortical regions was calculated and compared between brainstem nuclei. Results were aggregated to produce a freely available MNI structural atlas of limbic brainstem structures.

**Results:** A general trend was observed for a high probability of connectivity to the amygdala, hippocampus and DLPFC with relatively lower connectivity to the orbitofrontal cortex, nucleus accumbens, hippocampus and insula. The locus coeruleus and nucleus tractus solitarius demonstrated significantly greater connectivity to the DLPFC than amygdala while the periaqueductal grey, dorsal raphe nucleus, and ventral tegmental area did not demonstrate a significant difference between these two structures.

**Conclusion:** Monoaminergic and other modulatory nuclei in the brainstem project widely to cortical limbic regions. We describe the structural connectivity across the several key brainstem nuclei theorized to influence emotion, reward, and cognitive functions. An increased understanding of the anatomic basis of the brainstem’s role in emotion and other reward-related processing will support targeted neuromodulatary therapies aimed at alleviating the symptoms of neuropsychiatric disorders.

**Highlights:**

- The brainstem plays a key role in the processing of emotional stimuli and is intricately linked with the limbic system.
- Anatomic data for these connections is limited in humans
- We describe the structural connectivity of five brain stem nuclei (locus coeruleus, ventral tegmental area, periaqueductal grey, dorsal raphe nucleus, and nucleus tractus solitarius) in relation to limbic circuits
- Our results present a comprehensive delineation of the brainstem-limbic structural connectivity of these nuclei and are compiled into a freely available tractographic atlas
- Applications include future targeting of these structures for neuropsychiatric conditions

## Introduction

The brainstem, comprised of the medulla oblongata, pons, and midbrain, comprises approximately 3% of the mass of the brain and contains about 2% of the neurons in the central nervous system. Yet, what it lacks in size it makes up for in complexity and a disproportional influence on processes ranging from autonomic functions to arousal and consciousness.[1], [2] The brainstem provides vital autonomic regulation and homeostatic maintenance. It also serves as a conduit for all fibers linking the cerebral cortex and cerebellum to the spinal cord. Furthermore, it functions as a major afferent sensory system, receiving input from visceral fibers and cranial nerve nuclei which it then filters and transmits (often through several intermediary nuclei), to higher cortical centers.

The brainstem also plays a key role in emotional processing. Three brainstem networks have been identified that are thought to contribute to limbic processing: (1) the **ascending sensory network** consisting of the spinothalamic tracts, medial forebrain bundle, nucleus of the tractus solitarius (NTS), parabrachial nuclear complex and thalamic nuclei, (2) the **descending motor network** consisting of the periaqueductal grey (PAG), caudal raphe nucleus, and locus coeruleus (LC), and (3) the **modulatory network** with the serotonergic dorsal raphe (DRN), noradrenergic LC, and dopaminergic ventral tegmental area (VTA).[3], [4] The ascending, descending and modulatory brainstem networks allow for the progressive integration and processing of information as signals move rostrally through the brainstem, thalamus and then to the cortex, but also carry information in reverse, with cortical regions regulating the action-response relationships of phylogenetically older structures.[5]

The anatomic and structural basis for the brainstem’s role in limbic processing is theorized to involve several key nuclei which are the sole or major source of potent monoamine neurotransmitters for the higher cerebral cortex: the LC (norepinephrine), DRN (serotonin), and VTA (dopamine). Additionally, the PAG and NTS serve as inputs or centers of modulation to these monoamine neural networks (**Figure 1**). Several monoamine neurotransmitters have, individually or in combination, been implicated in disease states including Parkinson’s disease, major depressive disorder, or addiction, and are essential for physiological activities including arousal, sleep/wake cycles, perception of pain, affect, and goal directed behavior. Existing evidence obtained largely from animal studies **(Table 1)**, describes how these brainstem nodes interact with cortical limbic structures to convey body state and homeostatic information to result in behaviors such as heightened alertness, arousal from sleep, fear, and defense measures.[6] Overall, however, despite their importance, it has been challenging to study brainstem nuclei in humans in vivo because of difficulties in accurately defining the nuclei on imaging. Therefore, limited anatomic data exists for these brainstem-limbic relationship in humans.

**FIGURE 1:**
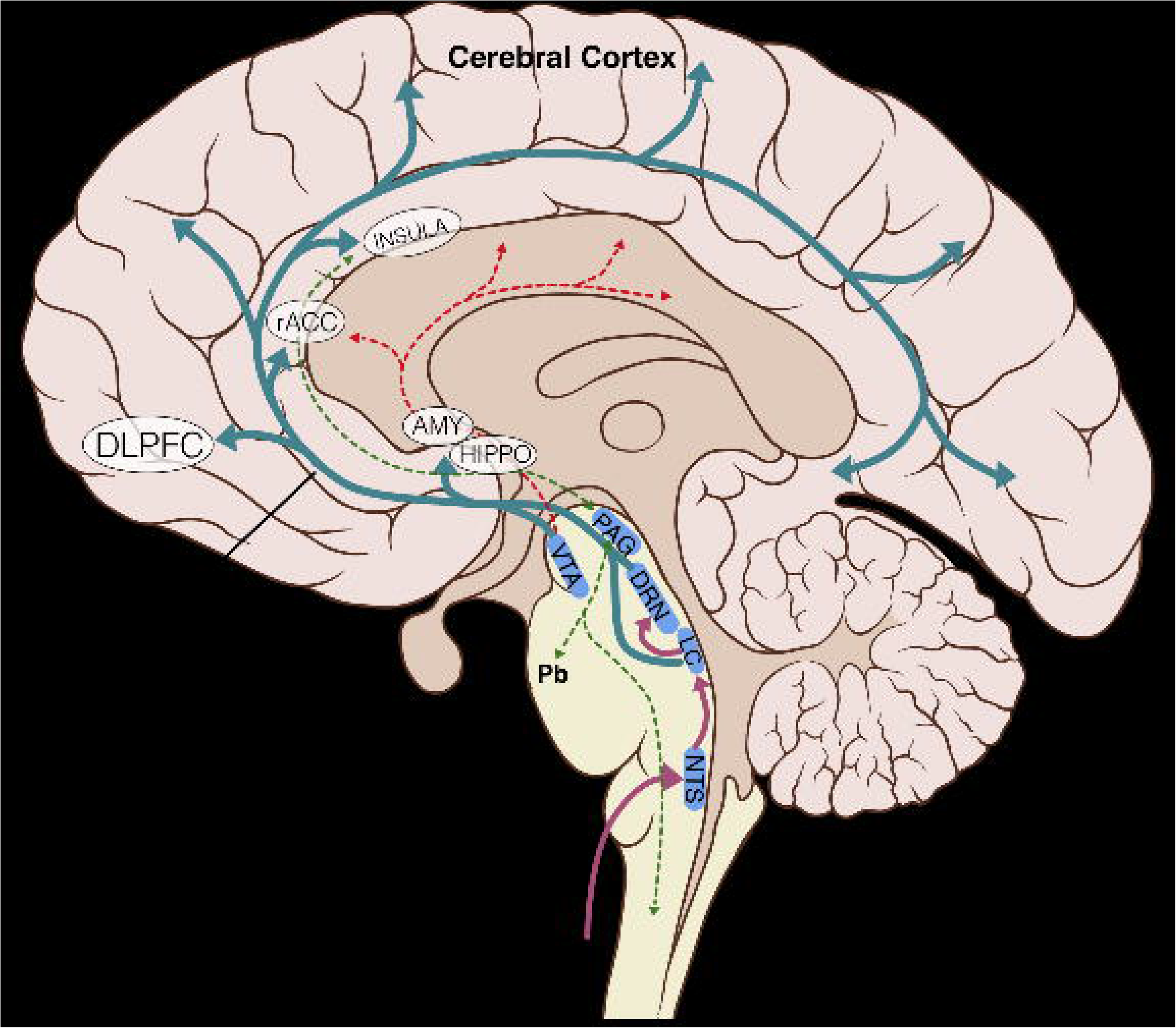
Schematic Representation of key brainstem-cortex circuits. DLPFC: dorsolateral-prefrontal cortex; rACC: rostral anterior cingulate cortex; INSULA: insular cortex; AMY: amygdala; HIPPO: hippocampus; VTA: ventral tegmental area; PAG: periaqueductal grey; DRN: dorsal raphe nucleus; LC: locus coeruleus; NTS: nucleus tractus solitarius; Pb: parabrachial nucleus; NE: norepinephrine; DA: dopamine; 5-HT: serotonin

**TABLE 1:**
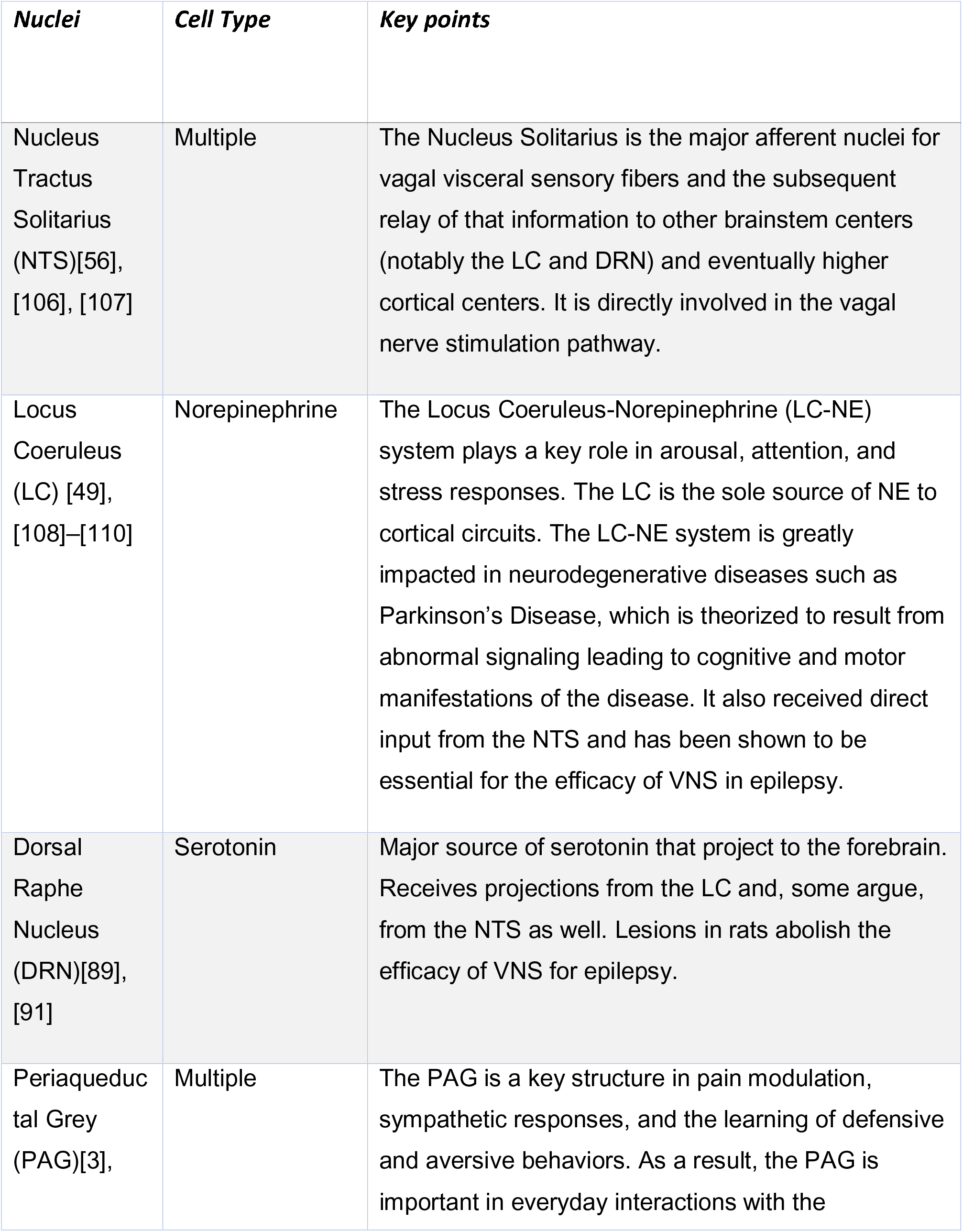
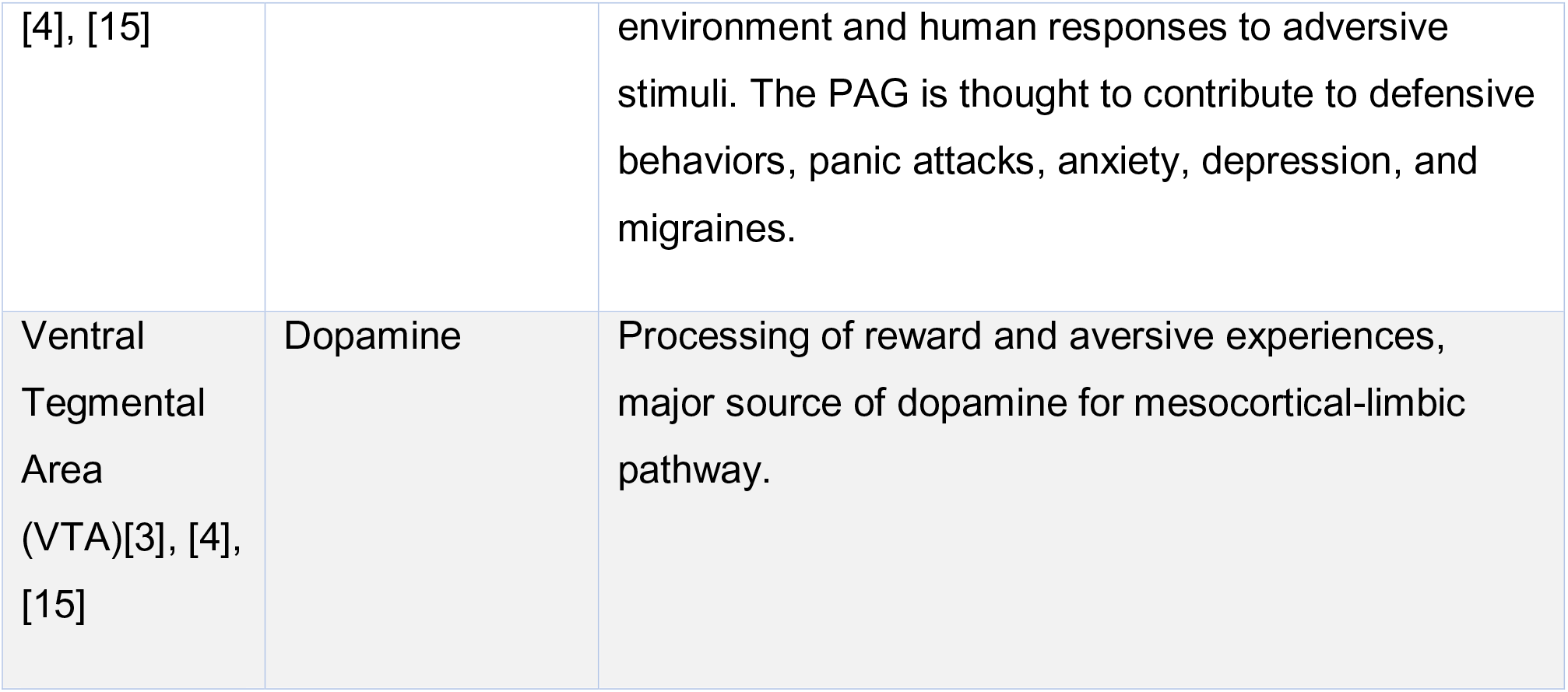
Limbic Brainstem Nuclei Key Characteristics.

There are two general approaches to defining regions of interest (ROI) on MRI, manual and automated segmentation. Automated segmentation has been demonstrated to reliably delineate cortical structures. [7], [8] It has also been used successfully to define certain regions of the brainstem (particularly for studies interested in masking the medulla, pons and midbrain separately), [9]–[12] yet it remains technically difficult to define most individual brainstem nuclei via this method. While there are several automated techniques, most recent attempts have been to mask brainstem nuclei using convolutional neural network-based segmentation, a deep learning image recognition technique, to delineate the substantia nigra [13] and LC. [14] However, this method relies on neuromelanin-MRI scans, limiting its application to nonpigmented regions. Furthermore, other techniques such as voxel intensity-based algorithms still require some manual delineation and thresholding and can be complicated by homogenously intense regions. [13]–[16] Therefore, although automated processing technologies are being developed, manual segmentation still has significant advantages and is considered the gold standard.

Manual segmentation, on the other hand, is labor intensive, often resulting in studies with small sample sizes, and high inter-rater variability. In addition, utilizing a manually-defined mask as an atlas for new subjects can be more computationally intensive than automated segmentation techniques. [13], [17] Yet, a well-trained individual, given proper anatomic knowledge and tools, can produce reliable results and manually delineated atlases are still considered by many to be the gold standard. [11], [17], [18]

In this study, we manually segmented five brainstem nuclei to perform probabilistic tractography to selected limbic targets in 197 human subjects. We hypothesized that autonomic and monoamine brainstem nuclei would demonstrate structural connectivity to cortical limbic regions, namely the amygdala, insula and hippocampus. Additionally, since there were no existing atlases of the structural connectivity of these nuclei, we aimed to produce an accurate anatomical atlas for use in future research.

We overcame some of the challenges of performing tractography on the brainstem by using a rigorous anatomic definition scheme with a combination of voxel measurements and anatomic landmarks to reduce variability while defining each nucleus. Additionally, we were able to achieve a scale of nearly 200 subjects, similar to many automated based approaches, thereby increasing the power of the study and reducing the effects of outliers or isolated errors. Lastly, we obtained diffusion MRI (dMRI) scans from the Human Connectome Project which provided a large dataset with scans acquired in a high resolution allowing for increased accuracy. [19], [20] Overall, our results present a comprehensive delineation of the brainstem-limbic structural connectivity of these nuclei and are compiled into a freely available tractographic atlas.

## Methods

### Subjects

Data were obtained from the publicly available WU-Minn HCP 1200 Subjects data release repository. [19], [20] The scanning protocol was approved by Human Research Protection Office (HRPO), Washington University (IRB# 201 204 036). No human subject experimental procedures were undertaken at the authors’ home institutions. The participants included in the HCP 1200 Subjects data release provided written informed consent as approved by the Washington University IRB. From this repository, 200 total non-twin subjects were randomly selected. The analysis was limited to these subjects based on available computational resources. Three subjects were excluded due to lack of required imaging data files. The remaining 197 subjects were included in our analyses and a description of their demographic characteristics is provided in **Table 2**.

**TABLE 2:**
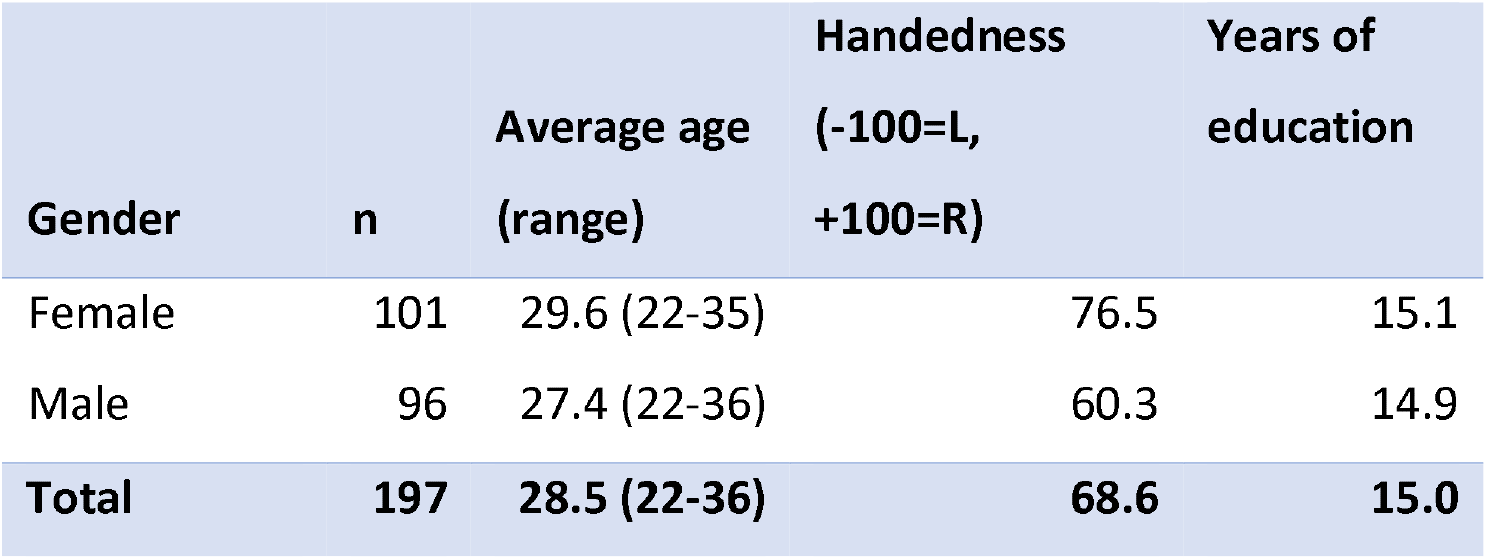
Cohort

| **Gender** | **n** | **Average age**<br>**(range)** | **Handedness**<br>**(-100=L,**<br>**+100=R)** | **Years of**<br>**education** |
| --- | --- | --- | --- | --- |
| Female | 101 | 29.6 (22-35) | 76.5 | 15.1 |
| Male | 96 | 27.4 (22-36) | 60.3 | 14.9 |
| **Total** | **197** | **28.5 (22-36)** | **68.6** | **15.0** |

### MRI Acquisition

The data were acquired on a modified Siemens 3T Skyra scanner with a customized protocol.^21^ The T1-weighted MRI has an isotropic spatial resolution of 0.7 mm, and the dMRI data have an isotropic spatial resolution of 1.25 mm. The multi-shell dMRI data were collected over 270 gradient directions distributed over three b-values (1000, 2000, 3000 s/mm2). For each subject, the multi-shell dMRI data were collected with both L/R and R/L phase encodings using the same gradient table, which were then merged into a single copy of multi-shell dMRI data after the correction of distortions with the HCP Preprocessing Pipeline. [20], [21]

### Masking of Seed Structures & Anatomic Boundaries

The areas of interest in this study consisted of brainstem structures with widespread projections. Due to the wide inter-subject anatomic variation and the small nature of many of these structures, all seed masks were created manually for each subject on the original Human Connectome Project (HCP) structural MRI scans using FSLeyes software. The seed masks generated in this fashion included the Locus Coeruleus (LC), Nucleus Tractus Solitarius (NTS), Periaqueductal Grey (PAG), Dorsal Raphe Nucleus (DRN) and Ventral Tegmental Area (VTA).

The anatomic boundaries used for each set of masks are described below and shown in **Figure 2**. Multiple histologic and radiographic resources were used to carefully define each region of interest.[22]–[26] Individual raters were first trained on a sample data set and their accuracy was assessed relative to a template mask. To ensure a high degree of internal consistency, no more than two individuals were responsible for creating the masks of each nucleus. After initial masking was completed, the fslmaths *boxv* command was used to dilate each mask by a factor of 5 followed by an erosion of 5 in order to ensure edges were smoothed and gaps were completely filled. The final version of each mask was independently verified by a neuroradiologist and neurosurgeon for anatomic accuracy before running tractography.

**FIGURE 2:**
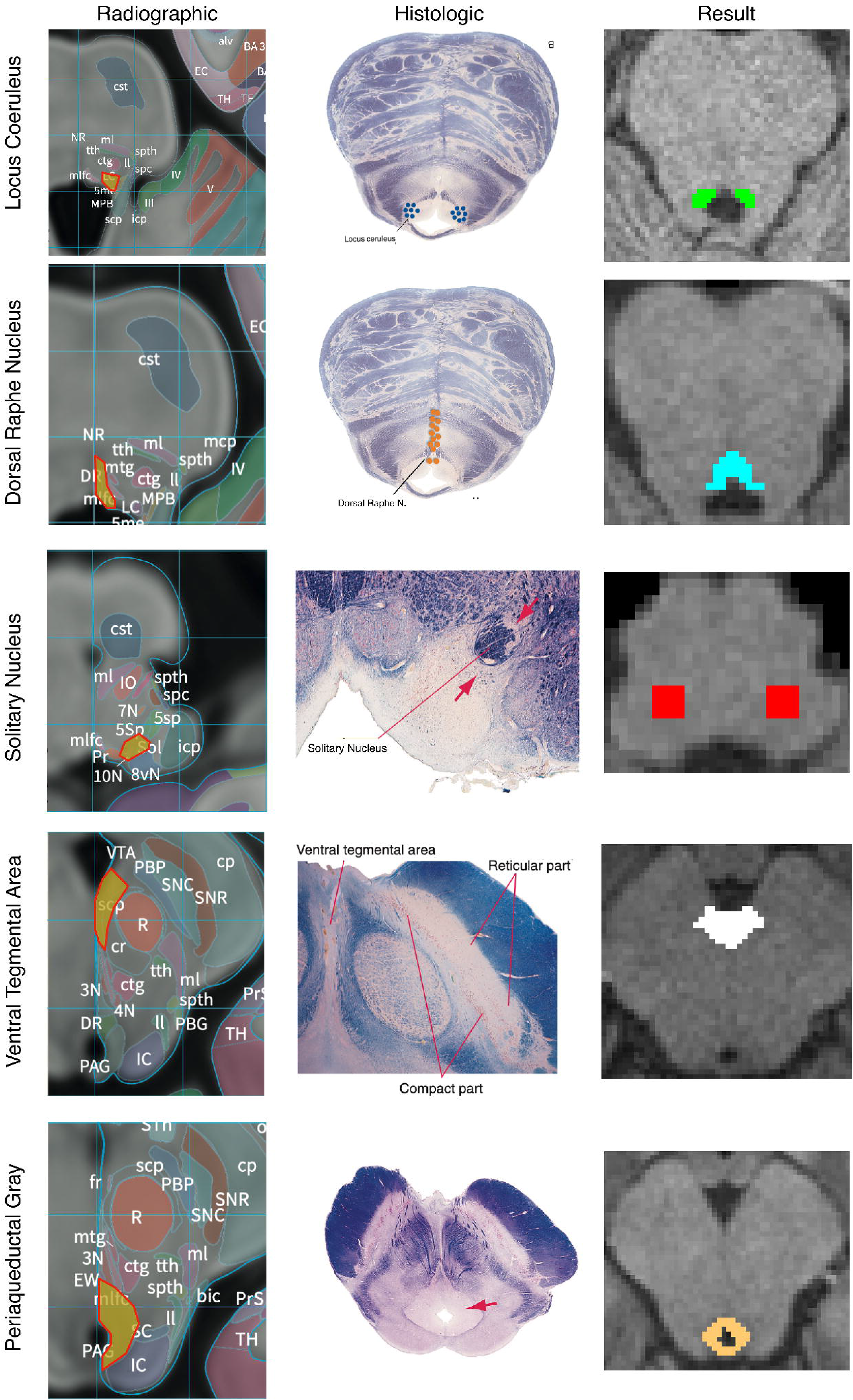
Anatomic Boundaries. Diagram of how anatomic boundaries of nuclei were masked. First radiographic^23^ (left pane) and histologic atlas^24^ (middle pane) were carefully examined for each nucleus. Using a combination of anatomic landmarks and voxel measurements (not shown), the resulting mask (right pane) was created in FSL.

### Locus Coeruleus

The LC was defined in both the caudal-rostral and medial-lateral planes. The most caudal point was the midpoint of the 4^th^ ventricle and the most rostral was the rostral pons. Laterally, the LC was defined to be 3mm lateral to the midline at the anterior-lateral angle of the 4^th^ ventricle (**FIGURE 2a**).

### Dorsal Raphe Nucleus

Beginning at the caudal midbrain, the DRN was masked caudally until its termination in the mid pons. The cerebral aqueduct and fourth ventricle were used as guides with the rostral portions measuring 12 voxels wide tapering to 6 voxels at the most caudal aspect (**FIGURE 2b**).

### Nucleus Tractus Solitarius

The caudal aspect of the NTS was defined to begin 3mm rostral to the obex. It then proceeded rostrally in a “V” shape. Standardized voxel measurements were used to ensure a consistent shape across individual. The NTS was masked caudally until the middle cerebellar peduncles were clearly visible on a horizontal section. Since the NTS also tracks slightly more anteriorly as it progresses rostrally, the anterior-caudal boundary was defined in the horizontal plane as the point where a line would transect from the root of cranial nerve VIII, with the point halfway between the midpoint and lateral aspect of the 4^th^ ventricle on either side of the pons (**FIGURE 2c)**.

### Ventral Tegmental Area

The most inferior transverse section of the VTA was defined at the section of the midbrain where the interpeduncular fossa opens to the interpeduncular cistern. The VTA boundary on inferior sections was the medial border of the Substantia Nigra. The lateral and medial borders were directly adjacent to the interpeduncular fossa. As the VTA progresses rostrally it becomes a contiguous structure with its medial boundaries joining in the midline bordering the medial aspect of the interpeduncular fossa and extending in a posterior direction to the midpoint of the medial edge of the red nucleus. The most superior section of the VTA was defined at the level of the mammillary bodies. Standardized voxel measurements were used to ensure a consistent shape across individual subjects (**FIGURE 2d**).

### Periaqueductal Grey

From a mid-coronal slice, the aqueduct was identified on the sagittal perspective. On the axial perspective, the lateral ventricles were traced inferiorly until it became the cerebral aqueduct which delineated the superior margin of the mask. A diamond-shaped border surrounding the aqueduct was demarcated as the PAG on the axial plane. The inferior border of the mask was determined to be the location where the mammillary bodies became fully defined, which correlated to the opening of the fourth ventricle. Standardized voxel measurements were used to ensure a consistent shape across individual subjects (**FIGURE 2e**).

### Probabilistic Tractography

Probabilistic tractography was carried out using FSL’s FMRIB Diffusion Toolbox (probtrackx) with modified Euler streaming. [7], [27] Target masks were generated using the Harvard-Oxford subcortical atlas.[25] Target masks included the amygdala (AMY), dorsolateral prefrontal cortex (DLPFC), hippocampus, insula, nucleus accumbens (NAc), orbitofrontal cortex (OFC), and rostral anterior cingulate cortex (rACC). Seed masks also served as target masks once produced such that the number of targets increased overtime as new seeds were created. All tractography was performed between each (right and left) seed and the ipsilateral target. Each target mask was also a termination mask such that tractography was terminated once a streamline entered the target. Additionally, Freesurfer standard lookup tables were used to generate Ipsilateral white matter masks which were used as waypoints. The ventricles and cerebellum masks were similarly generated with Freesufer and used as exclusion masks. We used the “onewaycondition”, curvature 0.2, 2000 samples, steplength=0.5, fibthresh=0.01, distthresh=1 and sampvox=0.0. This resulted in 14 or more seed_to_target output files representing a voxelwise map of the number of seed samples from each seed to target.

To calculate the **probability of connectivity (POC)** between each brainstem seed voxel to each of the 7 cortical and to the other 4 brainstem nuclei targets, we ran the FSL proj_thresh subroutine with a threshold of 1250 on each probtrackx output. For each voxel in the seed mask with a value above the threshold, *proj_thresh* calculates the number of samples reaching each of the target masks as a proportion of the total number of samples reaching any of the target masks. This yielded a separate map of each seed mask for each target with each voxel having a value between 0 and 1 representing the POC of that voxel to the given target. Thus, there were 2 maps (one for each hemisphere) per seed and per target for each subject. To produce an overall POC from each seed mask to target, probabilities were averaged across all voxels in each map. We next created a population connectivity map across all 197 subjects. Each of the previously created proj_thresh maps was registered to MNI 1mm standard space, thresholded at a level of 0.1 and binarized. These maps were then added across all 197 subjects such that each voxel value now represented the number of subjects with connectivity to the target.

### Parallel Data Processing

FSL software was implemented in a distributed fashion using Amazon Web Services (AWS, http://aws.amazon.com) EC2 instances running in parallel. Each AWS EC2 instance was an r4.large clone of an Amazon Machine Image (AMI) running Ubuntu 14.04 with FSL software version 5.0.10. FSL bedpostx directories for each subject and the probtrackx output files were stored on an Amazon S3 bucket.

### Statistical Analysis

Statistical analysis was carried out using the R software package (http://www.R-project.org/) and Prism 8 software from Graphpad (https://www.graphpad.com/).

To analyze the structural data, the subject specific output from the proj_thresh files stored on Amazon S3 were downloaded to a new EC2 instance running Ubuntu 14.04.1. FSLmaths was then used to compute the mean connectivity to each target using fslstats. The means were then imported to RStudio (version 1.3.959) running R (version 3.6.3) and the means to each target region were compared with a single factor analysis of variance (ANOVA). A Tukey HSD test was then run to determine statistical significance of the variance in the means.

Overall connectivity measurements were obtained by first taking the average of right and left connectivity for each subject specific seed-target relationship and then computing the mean across all subjects.

### MNI Atlas

Separately, each seed mask was warped to MNI space using the FSL *applywarp* command and the HCP subject specific nonlinear acpc_dc2standard file. The MNI152 NLIN 2009b T1 0.5mm brain was used as reference. [28] Once warped, each mask was averaged across all 197 subjects and set the threshold to two standard deviations from the mean to exclude extreme values.

## Results

Summative images in MNI space are publicly available at: https://www.uclahealth.org/neurosurgery/research-areas. Additionally, the atlas comes preinstalled on the widely used neurostimulation software, *Lead-DBS* (https://www.lead-dbs.org/helpsupport/knowledge-base/atlasesresources/atlases). All other data is available upon request of the corresponding author.

Complete results for the average POC for each seed to target relationship is reported in **TABLE 3**. **Figures 3-7** details the average streamline paths, provides a graphic representation of mean connectivity, and portrays the anatomic boundaries of each seed mask averaged over all subjects in MNI space. A summative three dimensional atlas is also showed (**FIGURE 8**). The three highest POCs are reported here for each seed.

**TABLE 3:** Average Probabilities of Connectivity; Format: Mean [95% CI interval]

|  | Dorsal Raphe Nucleus | Locus Coeruleus | Nucleus Tractus Solitarius | Periaqueductal Grey | Ventral Tegmental Area |
| --- | --- | --- | --- | --- | --- |
| *Amygdala* | 0.204 [0.195, 0.214] | 0.273 [0.262, 0.284] | 0.184 [0.176, 0.192] | 0.273 [0.262, 0.283] | 0.221 [0.210, 0.232] |
| *Dorsolateral Prefrontal Cortex* | 0.189 [0.179, 0.200] | 0.349 [0.333, 0.364] | 0.257 [0.244, 0.270] | 0.271 [0.257, 0.285] | 0.252 [0.236, 0.268] |
| *Hippocampus* | 0.124 [0.116, 0.131] | 0.150 [0.143, 0.157] | 0.108 [0.102, 0.113] | 0.157 [0.151, 0.164] | 0.174 [0.166, 0.182] |
| *Insula* | 0.077 [0.071, 0.082] | 0.110 [0.102, 0.117] | 0.089 [0.083, 0.095] | 0.145 [0.136, 0.154] | 0.083 [0.077, 0.089] |
| *Nucleus Accumbens* | 0.023 [0.021, 0.024] | 0.028 [0.026, 0.031] | 0.023 [0.021, 0.024] | 0.038 [0.035, 0.040] | 0.020 [0.018, 0.021] |
| *Orbital Prefrontal Cortex* | 0.070 [0.065, 0.074] | 0.095 [0.088, 0.102] | 0.067 [0.063, 0.072] | 0.106 [0.099, 0.112] | 0.071 [0.066, 0.076] |
| *Rostra Anterior Cingulate Cortex* | 0.010 [0.008, 0.011] | 0.009 [0.009, 0.010] | 0.010 [0.009, 0.011] | 0.020 [0.018, 0.022] | 0.006 [0.005, 0.006] |
| *Dorsal Raphe Nucleus* | - | - | - | - | 0.041 [0.036, 0.046] |
| *Locus Coeruleus* | 0.022 [0.020, 0.024] | - | 0.260 [0.244, 0.275] | 0.014 [0.011, 0.016] | 0.023 [0.021, 0.025] |
| *Nucleus Tractus Solitarius* | 0.123 [0.114, 0.132] | - | - | 0.035 [0.031, 0.039] | 0.140 [0.126, 0.154] |
| *Periaqueductal Grey* | 0.051 [0.046, 0.055] | - | 0.037 [0.031, 0.042] | - | 0.013 [0.012, 0.015] |
| *Ventral Tegmental Area* | 0.193 [0.177, 0.208] | - | - | - | - |

**FIGURE 3:**
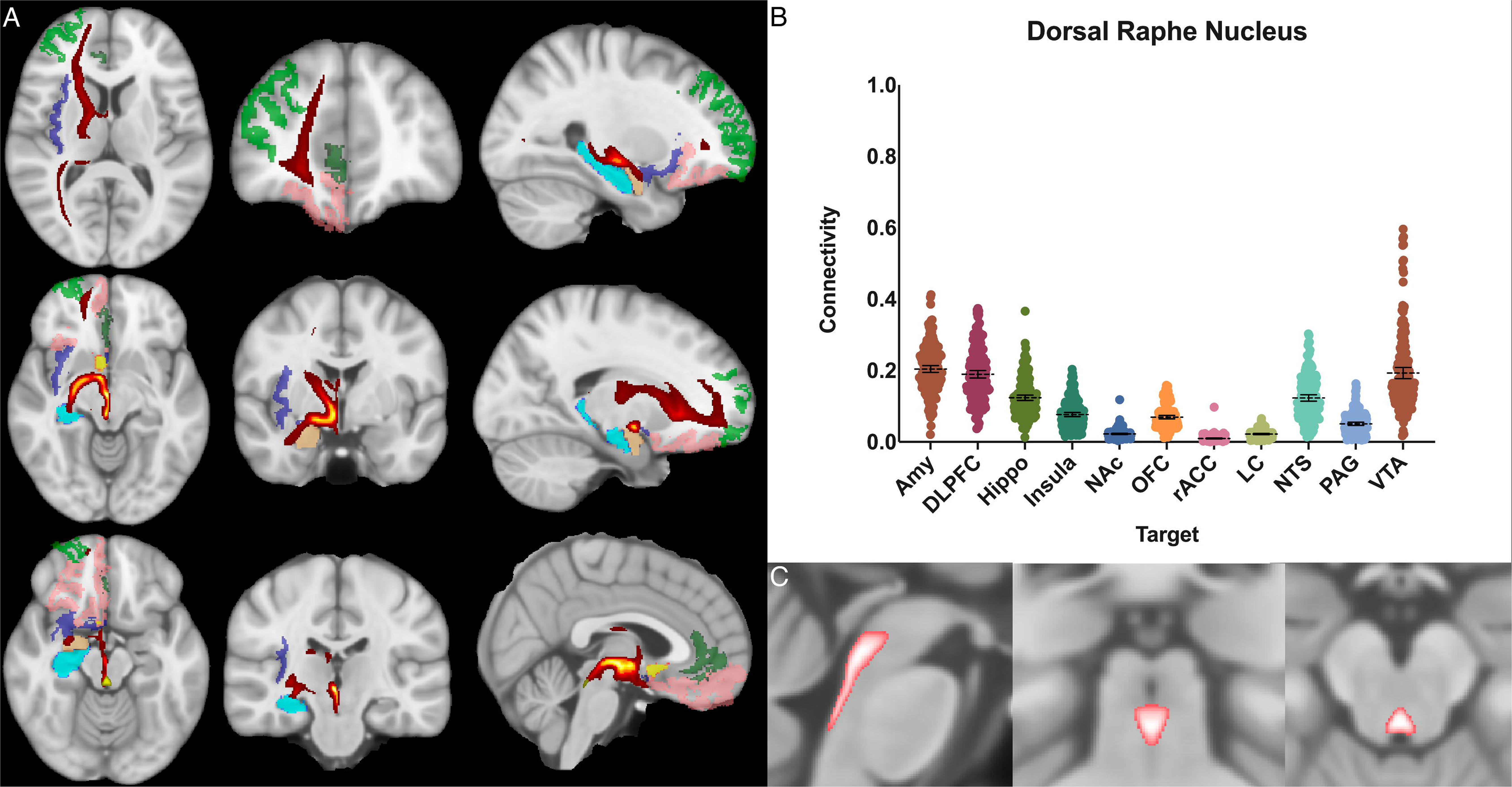
Dorsal Raphe Nucleus Structural Results. (A) MNI space structural connectivity results visual representation averaged over all 197 subjects. Heat map indicates Brighter yellow on heat map indicates a high number of samples passing through a given point that will eventually reach a target map (brighter yellow = more samples). Light green: DLPFC; dark green: OFC; light blue: AMY; brown: HIPPO; purple: insula; orange: nucleus accumbens; pink: rACC; yellow: seed (DRN). (B) Mean connectivity results with dashed line showing mean and 95% CI, each point on graph shows result from individual subject. (C) Anatomic MNI mask of seed region.

**FIGURE 4:**
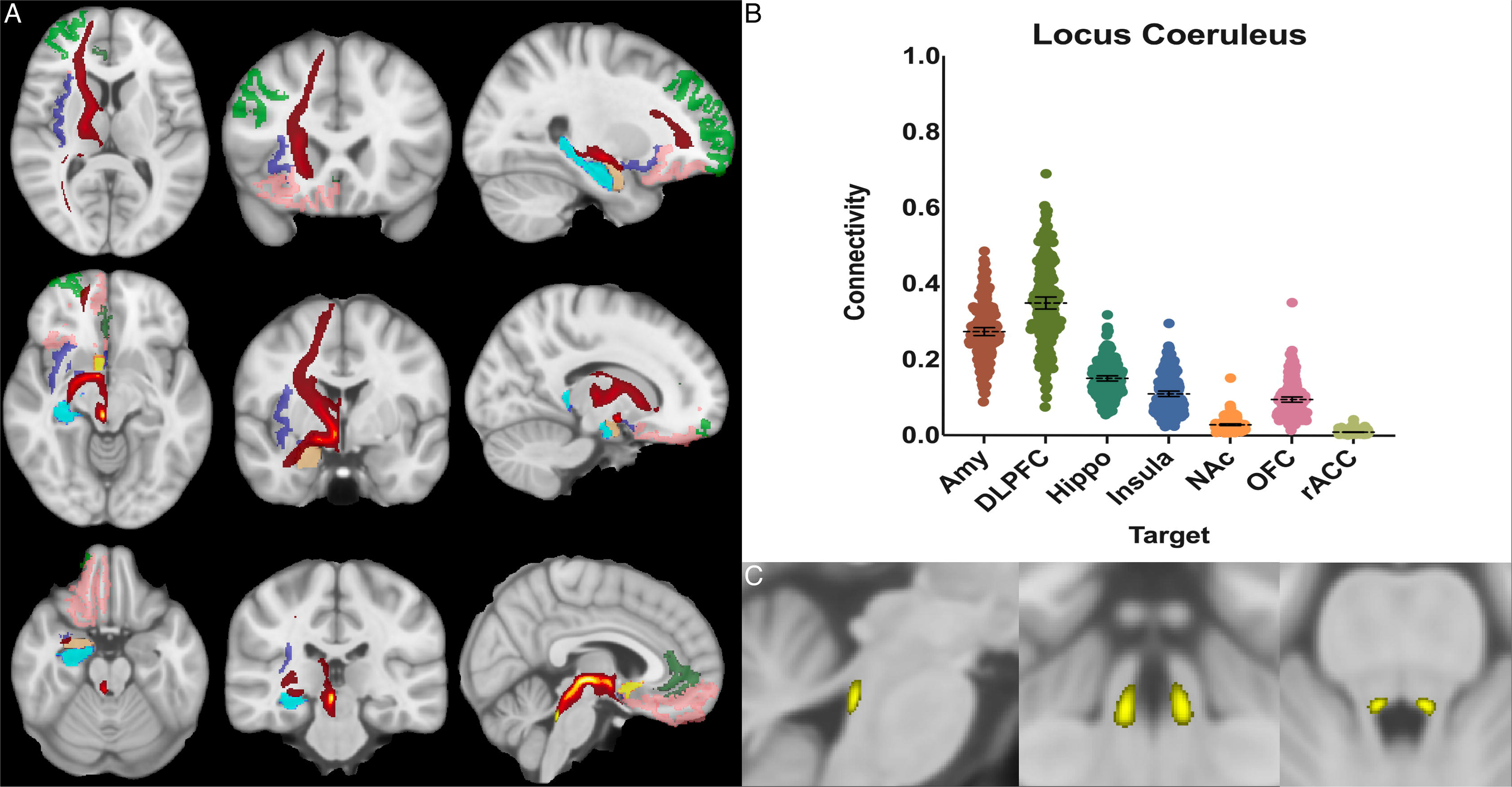
Locus Coeruleus Structural Results. (A) MNI space structural connectivity results visual representation averaged over all 197 subjects. Heat map indicates Brighter yellow on heat map indicates a high number of samples passing through a given point that will eventually reach a target map (brighter yellow = more samples). Light green: DLPFC; dark green: OFC; light blue: AMY; brown: HIPPO; purple: insula; orange: nucleus accumbens; pink: rACC; yellow: seed (LC). (B) Mean connectivity results with dashed line showing mean and 95% CI, each point on graph shows result from individual subject. (C) Anatomic MNI mask of seed region.

**FIGURE 5:**
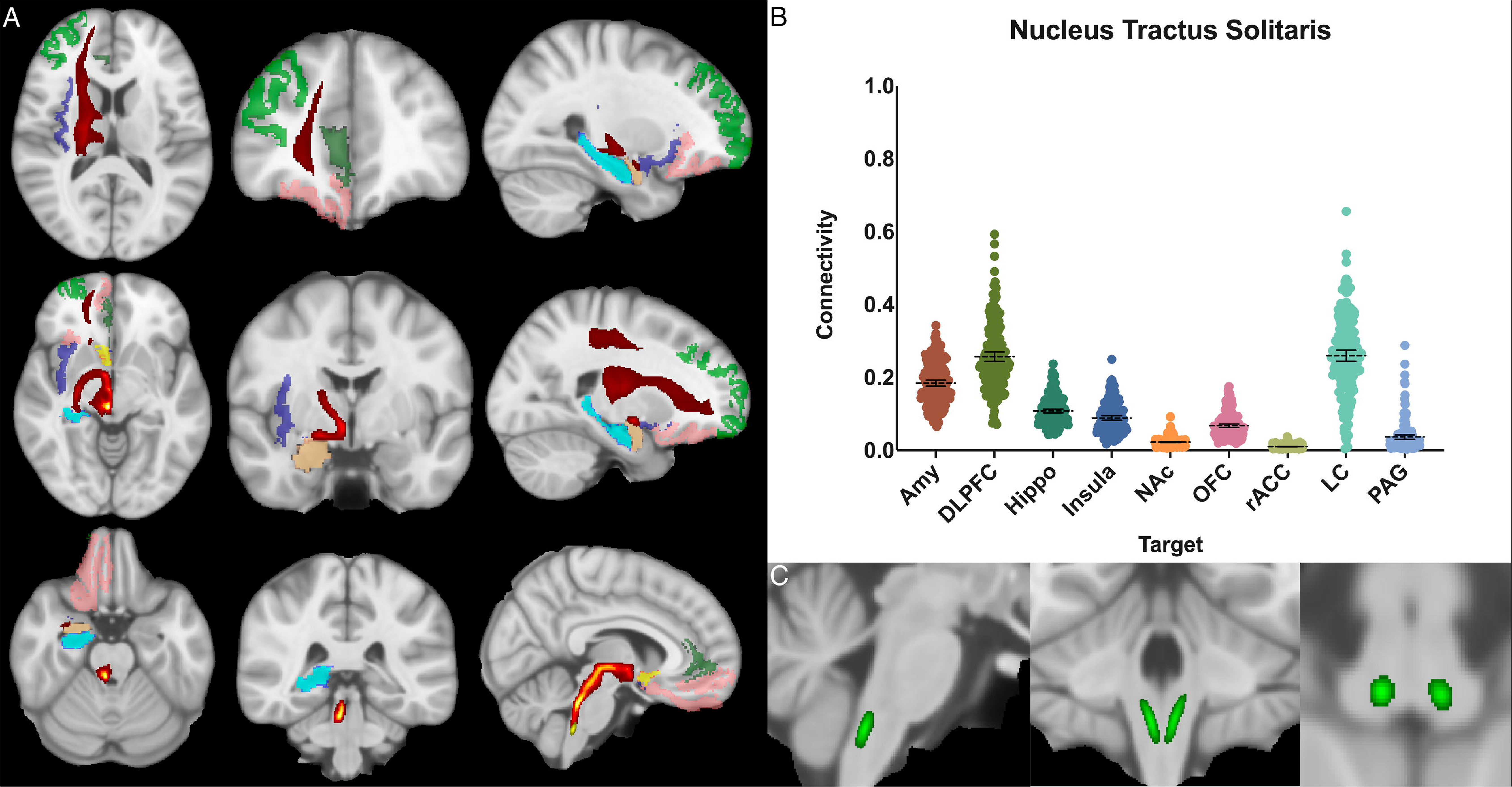
Nucleus Tractus Solitarius Structural Results. (A) MNI space structural connectivity results visual representation averaged over all 197 subjects. Heat map indicates Brighter yellow on heat map indicates a high number of samples passing through a given point that will eventually reach a target map (brighter yellow = more samples). Light green: DLPFC; dark green: OFC; light blue: AMY; brown: HIPPO; purple: insula; orange: nucleus accumbens; pink: rACC; yellow: seed (NTS). (B) Mean connectivity results with dashed line showing mean and 95% CI, each point on graph shows result from individual subject. (C) Anatomic MNI mask of seed region.

**FIGURE 6:**
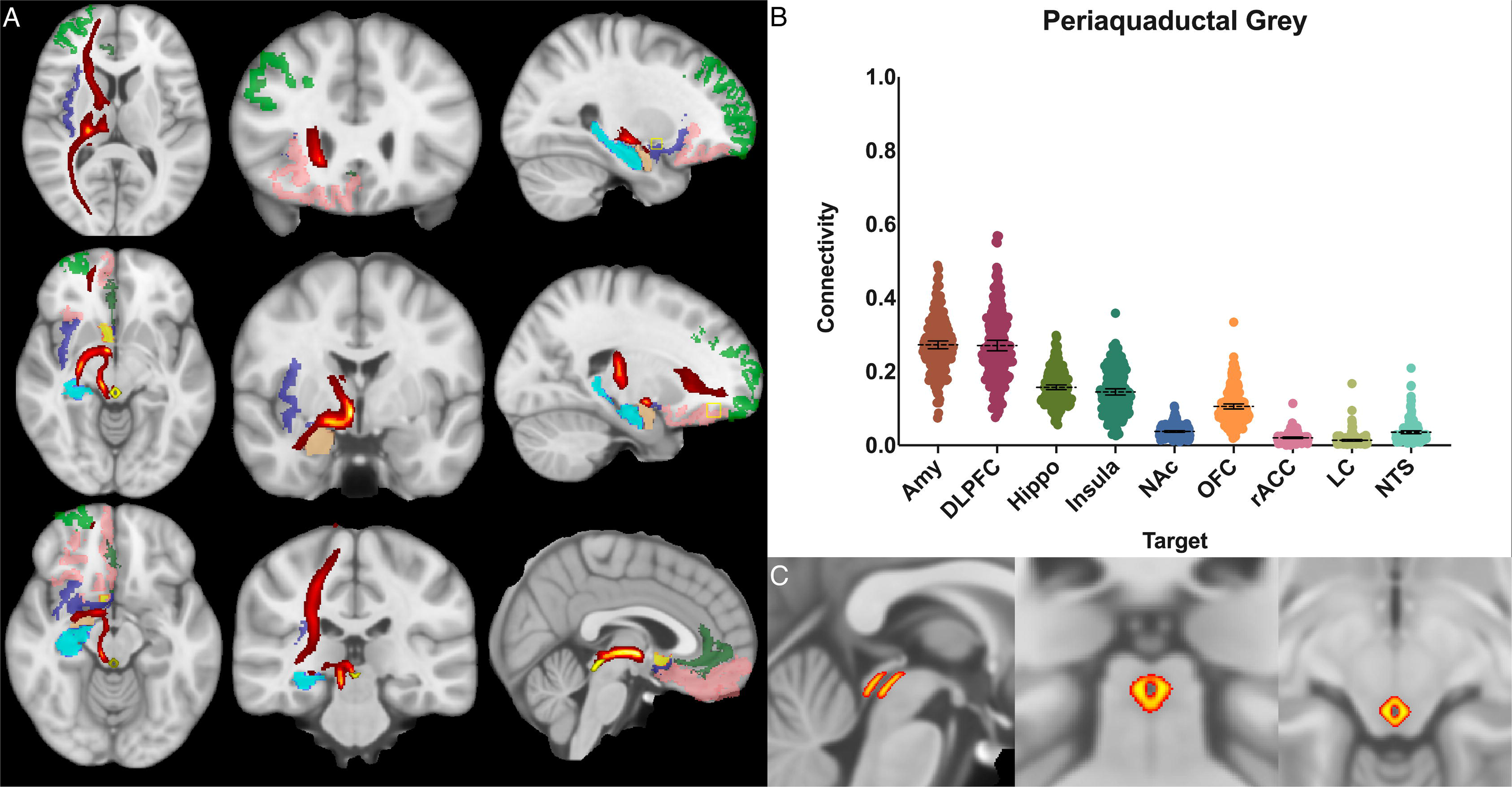
Ventral Tegmental Area Structural Connectivity. (A) MNI space structural connectivity results visual representation averaged over all 197 subjects. Heat map indicates Brighter yellow on heat map indicates a high number of samples passing through a given point that will eventually reach a target map (brighter yellow = more samples). Light green: DLPFC; dark green: OFC; light blue: AMY; brown: HIPPO; purple: insula; orange: nucleus accumbens; pink: rACC; yellow: seed (VTA). (B) Mean connectivity results with dashed line showing mean and 95% CI, each point on graph shows result from individual subject. (C) Anatomic MNI mask of seed region.

**FIGURE 7:**
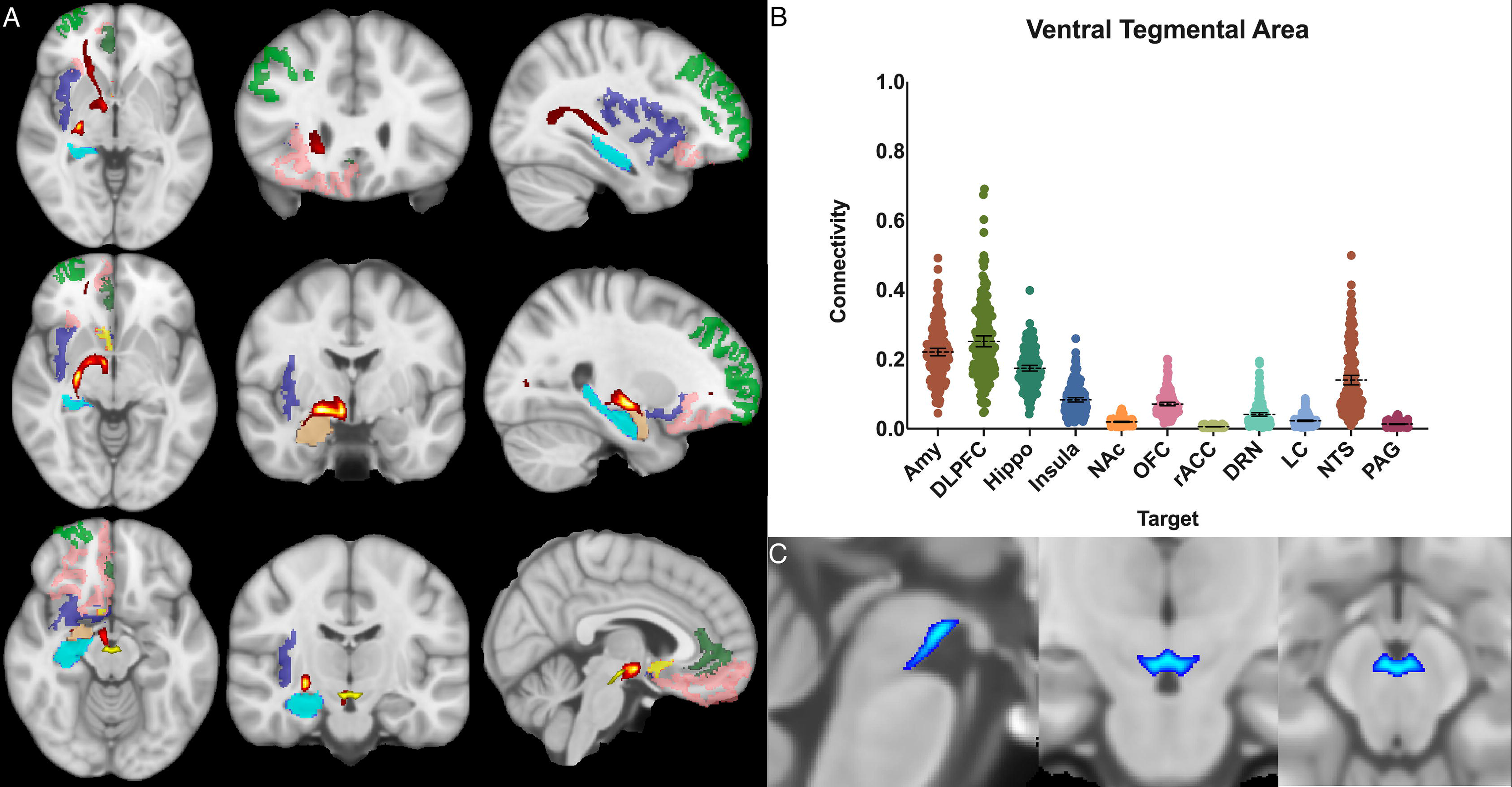
Periaqueductal Grey Structural Connectivity. (A) MNI space structural connectivity results visual representation averaged over all 197 subjects. Heat map indicates Brighter yellow on heat map indicates a high number of samples passing through a given point that will eventually reach a target map (brighter yellow = more samples). Light green: DLPFC; dark green: OFC; light blue: AMY; brown: HIPPO; purple: insula; orange: nucleus accumbens; pink: rACC; yellow: seed (PAG). (B) Mean connectivity results with dashed line showing mean and 95% CI, each point on graph shows result from individual subject. (C) Anatomic MNI mask of seed region.

**FIGURE 8:**
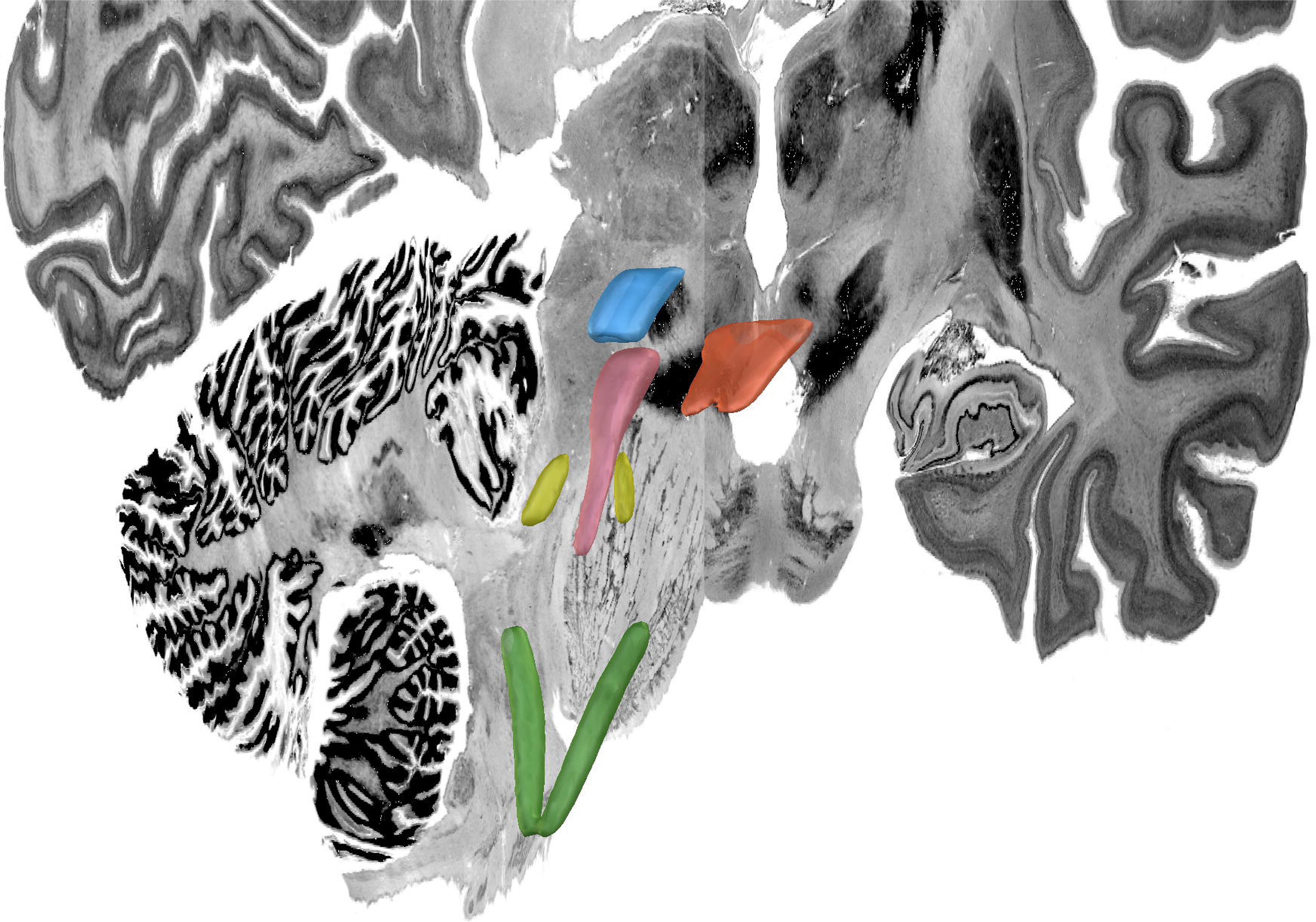
MNI 3D Atlas. Green: nucleus tractus solitaris; yellow: locus coeruleus; pink: periaqueductal grey; blue: ventral tegmental area; orange: dorsal raphe nucleus

Within each seed region, ANOVA with multiple comparisons (ANOVA-MC) and post-hoc t-test decompositions demonstrated significant (adjusted p<0.05) difference in the POC between each seed and target structure except as noted below. The only other exceptions to this were if the total POC was less than 0.03 for both structures.

The **dorsal raphe nucleus** demonstrated average POC to each target as follows: AMY 0.204 [95% CI 0.195, 0.214], VTA 0.193 [95% CI 0.177, 0.208], DLPFC 0.189 [95% CI 0.179, 0.200]. However, ANOVA-MC did not demonstrate a significant difference between these structures (**FIGURE 3.)**

The **locus coeruleus** demonstrated the greatest POC to the DLPFC 0.349 [95% CI 0.333, 0.364], followed by the AMY 0.273 [95% CI 0.262, 0.284] and Hippocampus 0.150 [95% CI 0.143, 0.157] (**FIGURE 4**).

The **nucleus tractus solitarius** showed the greatest POC to the LC 0.260 [95% CI 0.244, 0.275], followed by the DLPFC 0.257 [95% CI 0.244, 0.270] and the AMY 0.184 [95% CI 0.176, 0.192]. There was no significant difference between NTS-LC and NTS-DLPFC POC (**FIGURE 5**).

For the **periaqueductal grey** the highest POC was to the AMY 0.273 [95% CI 0.262, 0.283], followed by DLPFC 0.271 [95% CI 0.257, 0.285], however this difference was not significant (P>0.99). The third greatest POC was to the hippocampus at 0.157, significantly less than either the AMY (p<0.001) or the DLPFC (p<0.001) [95% CI 0.151, 0.164] (**FIGURE 6**).

The **ventral tegmental area** showed the greatest POC to the DLPFC 0.252 [95% CI 0.236, 0.268], AMY 0.221 [95% CI 0.210, 0.232], and the hippocampus 0.174 [95% CI 0.166, 0.182]. Of note, there was no significant difference between VTA-DLPFC and VTA-AMY POC (p=0.26) (**FIGURE 7**).

The demographic characteristics of the data set was homogenous by design **(TABLE 2)**. The t-test was performed between each seed-target POC and compared between males (n=96) and females (n=101). P-values were >0.05 for all comparisons with POC >0.03. The only exception was for the PAG-insula connection with male POC=0.135 and female POC=0.154 (P=0.025).

## Discussion

In this study, we carried out manual *in vivo* segmentation of defined brainstem regions of interest in a large cohort of subjects and performed probabilistic tractography between these regions and several limbic target regions of interest in the study of emotion and reward. The seeds selected were the NTS, LC, DRN, VTA, and PAG. These were chosen because of their clinical significance, the importance and wide-ranging effects of their respective neurotransmitters, and their established role in emotion and reward processing. We evaluated the connectivity between each of these seeds individually to a set of limbic processing centers to test the hypothesis of whether there would be a high POC between brainstem nuclei and these regions. Lastly, we aggregated these results into an MNI atlas and have made this publicly available to all interested researchers. This data represents a unique comprehensive *in vivo* analysis of key brainstem nuclei in a large human population and is also one of the first such studies to examine the relationship of these structures utilizing the Human Connectome Project.

Overall, our connectivity analysis of these five brainstem nuclei supports the role of the brainstem in emotional processing and confirmed our hypothesis that these brainstem nuclei demonstrate structural connectivity with known limbic regions. There is strong evidence that most mammals have analogous experiences to human emotion, even with far less evolved higher cortical centers, indicating that important processing occurs at levels below the cortex.[3], [4], [29], [30] In order to further understand the basis of emotion and reward, and intervene in cases of disorders of emotion and reward processing, it is critical to understand phylogenetically older structures such as the brainstem and recognize them as critical factors in emotional processing.

Emotions can be thought of as mental representations associated with distinct sensory states with the evolutionary goal of producing relevant behavioral responses in an organism. [3] It is therefore logical that the brainstem is a key modulator of this system as it is able to receive afferent visceral and somatic sensory information, begin filtering and processing these signals, and transmit them to higher cortical centers. Existing literature suggests that many of these processes are related to the monoamine neuropeptides, norepinephrine, serotonin, and dopamine. These neurotransmitters are produced by these brainstem nuclei which then project to higher cortical centers.[3], [22], [31], [32] Networks and nodes in the brainstem are increasingly considered key aspects of the conscious experience of emotion and therefore are of great clinical significance to neuropsychiatric disorders. [3], [22], [33]

In our analysis, each seed region demonstrated its own pattern of connectivity largely consistent with existing anatomical information from other imaging, postmortem and animal studies. [3], [24], [26], [33]–[35] A general trend was observed for high probabilities of connectivity with the amygdala, DLPFC, and hippocampus, with relatively lower connectivity with the OFC, NAc, insula, and rACC, suggesting that while there is heterogeneity among these nuclei, there is also a distinct pattern of brainstem connectivity with both frontal and temporal lobe structures.

### Dorsal Raphe Nucleus

The neurons which transmit serotonin and project to the cerebral cortex are mainly clustered around the dorsal and rostral aspects of the raphe nucleus. [36] The results here pertain only to the rostral portion as the largely inferiorly projecting caudal half was selectively excluded in the anatomic boundaries for this analysis (**FIGURE 2b**). The serotonin system is widely studied and is known to modulate fear and anxiety and other social behaviors. [37]–[40] Treatments for depression and anxiety uses medications selective serotonin reuptake inhibitors (SSRIs) targeted at this system.[41], [42] In animals, serotonergic projection to the amygdala and hippocampus has been associated with anxiety and retrieval of fear memories, and this relationship is under study in humans as a possible mechanistic explanation for major depressive disorder.[43]–[48]

Based on histological data, the DRN’s projection patterns are complex and include temporal lobe structures such as the hippocampus and amygdala, the PAG, LC, and frontal and insular cortices as well.[49] Functional connectivity studies have shown alterations in DRN connectivity to the frontal and cingulate cortices in subjects with depression.[46], [48] Given the DRN’s significance in depression, there have been studies analyzing DRN-amygdala structural connectivity.[42], [50] While, the DRN has been included in whole-brain brainstem connectivity studies which demonstrated a connectivity profile congruent with the histological evidence described above, [34] to our knowledge, this is the first study to focus specifically on the structural connectivity of the DRN to the limbic system in humans.

We found significant cortical connectivity of the DRN to the amygdala (0.20), DLPFC (0.19), hippocampus (0.12), insula (0.08) and OFC (0.07), as well as to brainstem structures such as the NTS (0.12) and VTA (0.19). There was also relatively greater variation between subjects for the DRN than for any other structure analyzed, congruent with prior work on the serotonin system that has shown significant difference across individuals. [51], [52] Interestingly, given the prior histologic evidence for LC-DRN and PAG-DRN connectivity, [49], [53], [54] we found a relatively low POC between these structures (0.02 and 0.05 respectively), which may reflect a limitation of our method for resolving extremely short-range connectivity within the brainstem (**FIGURE 3**). Taken together, our results provide important structural connectivity information, confirming significant connectivity between the DRN, amygdala, and hippocampus.

### Locus Coeruleus

Studies from nonhuman primates have shown that the LC receives viscerosensory inputs from the NTS and DRN, as well as descending information from the amygdala, OFC, and rACC. [55], [56] Recent work has also indicated that the LC plays a key role in shifting attention, [57], [58] emotionally salient memory formation and retrieval,[59]– [61] and cognition.[55], [56] Notable findings in our study confirmed strong connectivity to the NTS (0.25), hippocampus (0.15), and the amygdala (0.27), but showed the greatest overall connectivity between the LC and the DLPFC (0.35) (**FIGURE 4**). This was significantly greater than LC-AMY, LC-HIPPO, or LC-NTS (for all, p<0.0001) connectivity. This data supports electrophysiologic work in rats and non-human primates by demonstrating congruent anatomy in humans and supports the notion that a key role of the LC is to mediate attention, possibly through modulatory effects on the DLPFC, a cortical area known to become activated when subjects are asked to attend to specific stimuli.[55], [56] Furthermore, these findings highlight work that demonstrates the importance of the LC in mediating attention and sympathetic activation during acute stress and response to threats, systems which have been shown to be maladaptive in anxiety and depressive disorders.

### Nucleus Tractus Solitarius

The NTS is the major visceroafferent sensory nucleus for the vagal nerve complex.[53], [62] It receives approximately 75% of afferent visceral sensory information and relays this information to other nuclei within the brainstem, namely the LC and DRN.[54] While the NTS is involved in many physiologic functions including respiration and gastrointestinal regulation, it also has direct projections to the amygdala and has been implicated in panic disorder and memory formation.[59], [63] Prior imaging studies have sought to identify the NTS on high resolution imaging and perform tractography from it. [62], [64] However, differing anatomic definitions and the inherent difficulty of identifying the NTS on MRI, make meaningful comparisons challenging. Our data demonstrates significant connectivity between both the NTS and the amygdala (0.17) and NTS and the DLPFC (0.24), and we also show brainstem connections between the NTS and LC (0.26) and DRN (0.12) (**FIGURE 5**).

Given prior histological evidence, the expected findings of strong connectivity to the LC and DRN most likely indicate a pathway for afferent information from the vagus nerve to ascend to higher cortical centers.[54], [65], [66] Interestingly, we also found a relatively high POC to the amygdala and the DLPFC. Studies in rodents have indicated that the NTS may be involved in fear memory formation via its connections with the amygdala. [59], [63] Additional evidence supports that vagus nerve stimulation improves memory consolidation, and over 75% of vagal afferents project to the NTS. [23], [53], [67] Our results provide further evidence to support the anatomic basis of these findings in humans and could be used as the basis for further study of these phenomenon utilizing this atlas.

Clinically, the data on the NTS, DRN, and LC are interesting to consider in the context of existing work on vagus nerve stimulation (VNS). The NTS is thought to be the main vagal afferent nucleus for “body state information” (visceral sensory information from cardiopulmonary and digestive system). [54], [65], [66], [68]–[72] Based on animal and tract tracing studies, a large portion of fibers subsequently project to the LC and DRN which send adrenergic and serotonergic projections to cortical structures. [3]–[5] While it is not possible to delineate specific fiber types from this analysis, we found high POC between the LC-NTS (0.26), NTS-DRN (0.12), LC-AMY (0.27), and DRN-AMY (0.20). (**FIGURE 3-5**). Both the LC and DRN showed comparably strong connectivity to the DLPFC, Hippo, and Insula as well. Taken together, this atlas could prove useful to study the mechanism of VNS and possible structural reorganization of these pathways after a therapeutic intervention.

### Ventral Tegmental Area

The three main midbrain regions containing dopaminergic neurons are the retrorubral field, substantia nigra pars compacta, and VTA. The main dopaminergic system involved in limbic processes is the mesocorticolimbic pathway (**Figure 1),** which arises from the VTA and projects via the medial forebrain bundle to the nucleus accumbens and prefrontal cortex. [73]–[77] The VTA also receives feedback inhibition from the striatum, cortex, as well as the hippocampus and amygdala.[73]–[75], [78]–[80] The VTA is involved in motivation, reward, and arousal, and the dopamine system is known to be affected in disease states such as Parkinson’s Disease, addiction, depression, and schizophrenia. [3], [46], [76], [77], [81]

In our analysis, the greatest POCs were between the VTA and the DLPFC (0.25), amygdala (0.22), hippocampus (0.17) and NTS (0.14). (**FIGURE 6)**. Interestingly, despite the well-established relationship between the VTA and the nucleus accumbens, we found low POC (0.02) between these two structures. This is likely because our methodology compares relative POC between target regions and, though we do control for it, can still be affected by the total number of fibers between a seed and target. Our findings regarding VTA-DLPFC connectivity and VTA-AMY connectivity are particularly notable as several DTI studies have previously linked the VTA with frontal lobe areas (namely the prefrontal cortex and OFC) but did not assess or demonstrate significant connectivity with temporal lobe structures (namely the amygdala and hippocampus).[82]–[85] Here, we found no significant difference between VTA-DLPFC or VTA-AMY POC (p=0.26). This, combined with relatively low OFC POC (0.07) and a relatively high hippocampus POC (0.17), demonstrates a distinct connectivity pattern to *both* frontal and temporal lobe structures. The VTA is a potential target for deep brain stimulation as well as other neuromodulation techniques, and it is therefore highly important to consider its connectivity profile.[76], [86], [87] While dopaminergic connections between the VTA, amygdala, and hippocampus have previously been established, much of what is known is based on animal studies, and demonstrating this consistently in humans is of vital importance for the development of therapies based on this anatomy.[73], [77]

### Periaqueductal Grey

The PAG is a complex, heterogenous group of neurons that interacts with many brain regions and has roles in cardiorespiratory control, pain, fear, anxiety, and goal directed defensive behaviors.[88]–[90] Prior DTI studies have demonstrated PAG and prefrontal cortex connectivity. This is of interest because this connection has been suggested as a mechanism for the conscious modulation of pain signals.[89]–[93] Furthermore, amygdala and insular connections have been demonstrated and are theorized to be involved in the emotional response to pain. [90], [92]–[95]

We find a nonselective pattern of connectivity with POC values above 0.10 for the OFC, insula, hippocampus, DLPFC and amygdala, confirming both the frontal and temporal connectivity patterns found in previous studies described above. The only brainstem region with significant connectivity was the DRN (0.10), consistent with the theory that these two regions encircling the aqueduct are closely linked and involved in the processing of aversive stimuli (**FIGURE 7**).[43], [89], [91] While much of the data generated here was previously known, we sought to include the PAG in our atlas given its interplay with the DRN and importance in chronic pain.

### Modulation

Several attempts have been made to modulate these brainstem regions with varying degrees of success. [86], [87], [96]–[98] By providing here an atlas from a large cohort of subjects, we hope this data can be potentially used to help target brainstem nuclei more effectively (and/or their upstream or downstream targets) in future trials of deep brain stimulation for neuropsychiatric disorders.

## Limitations

There are several important limitations to this study. There are inherent limitations described in detail regarding registration and tractography.[99]–[102] The high-quality imaging protocols of the Human Connectome Project address several of the concerns related to artifact and scan acquisition. [19] Additionally, the large scale of our study means that artifactual errors in individual scans have a minimal (<1%) impact on our results. Another limitation involves potential errors in anatomic masking. Care must be taken to accurately mask these brainstem structures as small anatomic errors, especially at early stages, can be compounded in the analysis. We attempted to address this by carefully creating each subject specific mask by hand (**FIGURE 2**), having multiple individuals including a neuroradiologist independently check each mask for accuracy, by using high quality scans and reference resources, and by clearly defining our anatomic boundaries in this paper. Our methods allowed for resolution down to the size of individual voxels for each subject specific seed mask. A third limitation was that we did not run whole brain tractography and may be missing fiber tracts that did not involve our preselected target regions. However, as we were interested specifically in limbic connectivity, especially for monoamine nuclei, this method allowed us to appreciate differences between specific regions of interest more clearly than whole brain tractography. Additionally, since it is likely that some of the seed masks overlapped with large white matter tracts that abut these structures, this method prevented large motor and sensory tracts from complicating the analysis. Lastly, our method cannot determine directionality and care should be taken in ascribing any directional connectivity.

## Future Directions

Given that the Human Connectome Project has a significant amount of subject level behavioral and behavioral data available, an extension of this project currently underway is to correlate structural connectivity with emotion-related behavioral traits. Future work is needed to elucidate if structural variation in individuals is associated with behavioral or personality traits.

## Conclusion

The brainstem is an essential component of the limbic system. Monoamine and other modulatory nuclei in the brainstem project widely to cortical and subcortical limbic regions and each has a specific pattern of connectivity. An understanding of this basic structural anatomy is a critical step in understanding disease processes, such as addiction, chronic pain, and depression and the development of novel therapeutics. Further studies are warranted to characterize the functional significance of the structural connectivity of each nucleus and the relationship of structural connectivity with neuropsychiatric symptoms.

## Acknowledgement

A.H. was supported by the German Research Foundation (Deutsche Forschungsgemeinschaft, Emmy Noether Stipend 410169619 and 424778381 – TRR 295), Deutsches Zentrum für Luft- und Raumfahrt (DynaSti grant within the EU Joint Programme Neurodegenerative Disease Research, JPND), the National Institutes of Health (2R01 MH113929) as well as the Foundation for OCD Research (FFOR).

## References

[1] F. A. C. Azevedo et al., “Equal numbers of neuronal and nonneuronal cells make the human brain an isometrically scaled-up primate brain,” J. Comp. Neurol., vol. 513, no. 5, pp. 532–541, Apr. 2009, doi: 10.1002/cne.21974.

[2] S. Herculano-Houzel, “The human brain in numbers: a linearly scaled-up primate brain,” Front. Hum. Neurosci., vol. 3, 2009, doi: 10.3389/neuro.09.031.2009.

[3] A. Venkatraman, B. L. Edlow, and M. H. Immordino-Yang, “The Brainstem in Emotion: A Review,” Front. Neuroanat., vol. 11, 2017, doi: 10.3389/fnana.2017.00015.

[4] M. Angeles Fernández-Gil, R. Palacios-Bote, M. Leo-Barahona, and J. P. Mora-Encinas, “Anatomy of the brainstem: a gaze into the stem of life,” Semin. Ultrasound CT MR, vol. 31, no. 3, pp. 196–219, Jun. 2010, doi: 10.1053/j.sult.2010.03.006.

[5] D. M. Tucker, D. Derryberry, and N. O. Emotion, Vertical Integration of Brainstem, Limbic, and Cortical Systems. In J. Borod (Ed.), Handbook of the Neuropsychology of Emotion. New York: Oxford.

[6] Y. M. Ulrich-Lai and J. P. Herman, “Neural regulation of endocrine and autonomic stress responses,” Nat Rev Neurosci, vol. 10, no. 6, pp. 397–409, Jun. 2009, doi: 10.1038/nrn2647.

[7] M. W. Woolrich et al., “Bayesian analysis of neuroimaging data in FSL,” Neuroimage, vol. 45, no. 1 Suppl, pp. S173–186, Mar. 2009, doi: 10.1016/j.neuroimage.2008.10.055.

[8] B. Fischl et al., “Whole brain segmentation: automated labeling of neuroanatomical structures in the human brain,” Neuron, vol. 33, no. 3, pp. 341–355, Jan. 2002, doi: 10.1016/s0896-6273(02)00569-x.

[9] L. Sander et al., “Accurate, rapid and reliable, fully automated MRI brainstem segmentation for application in multiple sclerosis and neurodegenerative diseases,” Human Brain Mapping, vol. 40, no. 14, pp. 4091–4104, 2019, doi: https://doi.org/10.1002/hbm.24687.

[10] J. Y. Wang, M. M. Ngo, D. Hessl, R. J. Hagerman, and S. M. Rivera, “Robust Machine Learning-Based Correction on Automatic Segmentation of the Cerebellum and Brainstem,” PLOS ONE, vol. 11, no. 5, p. e0156123, May 2016, doi: 10.1371/journal.pone.0156123.

[11] J. E. Iglesias et al., “Bayesian segmentation of brainstem structures in MRI,” Neuroimage, vol. 113, pp. 184–195, Jun. 2015, doi: 10.1016/j.neuroimage.2015.02.065.

[12] B. Patenaude, S. M. Smith, D. N. Kennedy, and M. Jenkinson, “A Bayesian model of shape and appearance for subcortical brain segmentation,” Neuroimage, vol. 56, no. 3, pp. 907–922, Jun. 2011, doi: 10.1016/j.neuroimage.2011.02.046.

[13] A. Le Berre et al., “Convolutional neural network-based segmentation can help in assessing the substantia nigra in neuromelanin MRI,” Neuroradiology, vol. 61, no. 12, pp. 1387–1395, Dec. 2019, doi: 10.1007/s00234-019-02279-w.

[14] M. Dünnwald, M. J. Betts, A. Sciarra, E. Düzel, and S. Oeltze-Jafra, “Automated Segmentation of the Locus Coeruleus from Neuromelanin-Sensitive 3T MRI Using Deep Convolutional Neural Networks,” in Bildverarbeitung für die Medizin 2020, Wiesbaden, 2020, pp. 61–66. doi: 10.1007/978-3-658-29267-6_13.

[15] K. Ogisu et al., “3D neuromelanin-sensitive magnetic resonance imaging with semi-automated volume measurement of the substantia nigra pars compacta for diagnosis of Parkinson’s disease,” Neuroradiology, vol. 55, no. 6, pp. 719–724, Jun. 2013, doi: 10.1007/s00234-013-1171-8.

[16] S. Zhang, S. Hu, H. H. Chao, and C. S. Li, “Resting-State Functional Connectivity of the Locus Coeruleus in Humans: In Comparison with the Ventral Tegmental Area/Substantia Nigra Pars Compacta and the Effects of Age,” Cereb Cortex, vol. 26, no. 8, pp. 3413–27, Aug. 2016, doi: 10.1093/cercor/bhv172.

[17] P. Aljabar, R. A. Heckemann, A. Hammers, J. V. Hajnal, and D. Rueckert, “Multi-atlas based segmentation of brain images: atlas selection and its effect on accuracy,” Neuroimage, vol. 46, no. 3, pp. 726–738, Jul. 2009, doi: 10.1016/j.neuroimage.2009.02.018.

[18] R. A. Morey et al., “A comparison of automated segmentation and manual tracing for quantifying hippocampal and amygdala volumes,” Neuroimage, vol. 45, no. 3, pp. 855–866, Apr. 2009, doi: 10.1016/j.neuroimage.2008.12.033.

[19] D. C. Van Essen, S. M. Smith, D. M. Barch, T. E. J. Behrens, E. Yacoub, and K. Ugurbil, “The WU-Minn Human Connectome Project: An Overview,” Neuroimage, vol. 80, pp. 62–79, Oct. 2013, doi: 10.1016/j.neuroimage.2013.05.041.

[20] M. F. Glasser et al., “The minimal preprocessing pipelines for the Human Connectome Project,” Neuroimage, vol. 80, pp. 105–124, Oct. 2013, doi: 10.1016/j.neuroimage.2013.04.127.

[21] S. N. Sotiropoulos et al., “Advances in diffusion MRI acquisition and processing in the Human Connectome Project,” Neuroimage, vol. 80, pp. 125–143, Oct. 2013, doi: 10.1016/j.neuroimage.2013.05.057.

[22] B. L. Edlow et al., “Neuroanatomic connectivity of the human ascending arousal system critical to consciousness and its disorders,” J. Neuropathol. Exp. Neurol., vol. 71, no. 6, pp. 531–546, Jun. 2012, doi: 10.1097/NEN.0b013e3182588293.

[23] T. W. Vanderah, Nolte’s the human brain in photographs and diagrams, 5th edition. Philadelphia, PA: Elsevier, 2019.

[24] J. K. Mai, M. Majtanik, and G. Paxinos, Atlas of the human brain. 2016.

[25] R. S. Desikan et al., “An automated labeling system for subdividing the human cerebral cortex on MRI scans into gyral based regions of interest,” Neuroimage, vol. 31, no. 3, pp. 968–980, Jul. 2006, doi: 10.1016/j.neuroimage.2006.01.021.

[26] T. P. Naidich, H. M. Duvernoy, B. N. Delman, A. G. Sorensen, S. S. Kollias, and E. M. Haacke, Duvernoy’s Atlas of the Human Brain Stem and Cerebellum: High-Field MRI, Surface Anatomy, Internal Structure, Vascularization and 3 D Sectional Anatomy. Springer Science & Business Media, 2009.

[27] M. Jenkinson, C. F. Beckmann, T. E. J. Behrens, M. W. Woolrich, and S. M. Smith, “FSL,” Neuroimage, vol. 62, no. 2, pp. 782–790, Aug. 2012, doi: 10.1016/j.neuroimage.2011.09.015.

[28] “ICBM 152 Nonlinear atlases (2009) – NIST.” http://nist.mni.mcgill.ca/?p=904 (accessed Mar. 04, 2021).

[29] G. Holstege, “How the Emotional Motor System Controls the Pelvic Organs,” Sexual Medicine Reviews, vol. 4, no. 4, pp. 303–328, Oct. 2016, doi: 10.1016/j.sxmr.2016.04.002.

[30] A. D. Craig, “Interoception: the sense of the physiological condition of the body,” Curr. Opin. Neurobiol., vol. 13, no. 4, pp. 500–505, Aug. 2003, doi: 10.1016/s0959-4388(03)00090-4.

[31] G. Holstege, “The emotional motor system,” Eur J Morphol, vol. 30, no. 1, pp. 67– 79, 1992.

[32] R. Plutchik, “The Nature of Emotions: Human emotions have deep evolutionary roots, a fact that may explain their complexity and provide tools for clinical practice,” American Scientist, vol. 89, no. 4, pp. 344–350, 2001.

[33] B. L. Edlow, J. A. McNab, T. Witzel, and H. C. Kinney, “The Structural Connectome of the Human Central Homeostatic Network,” Brain Connectivity, vol. 6, no. 3, pp. 187–200, Nov. 2015, doi: 10.1089/brain.2015.0378.

[34] M. Bianciardi et al., “In vivo functional connectome of human brainstem nuclei of the ascending arousal, autonomic, and motor systems by high spatial resolution 7-Tesla fMRI,” MAGMA, vol. 29, no. 3, pp. 451–462, Jun. 2016, doi: 10.1007/s10334-016-0546-3.

[35] Y. Tang, W. Sun, A. W. Toga, J. M. Ringman, and Y. Shi, “A probabilistic atlas of human brainstem pathways based on connectome imaging data,” Neuroimage, vol. 169, pp. 227–239, Apr. 2018, doi: 10.1016/j.neuroimage.2017.12.042.

[36] J.-P. Hornung and N. De Tribolet, “Chemical Organization of the Human Cerebral Cortex,” in Neurotransmitters in the Human Brain, D. J. Tracey, G. Paxinos, and J. Stone, Eds. Boston, MA: Springer US, 1995, pp. 41–60. doi: 10.1007/978-1-4615-1853-2_4.

[37] H. Zangrossi, M. B. Viana, J. Zanoveli, C. Bueno, R. L. Nogueira, and F. G. Graeff, “Serotonergic regulation of inhibitory avoidance and one-way escape in the rat elevated T-maze,” Neuroscience & Biobehavioral Reviews, vol. 25, no. 7, pp. 637–645, Dec. 2001, doi: 10.1016/S0149-7634(01)00047-1.

[38] D. S. Moskowitz, G. Pinard, D. C. Zuroff, L. Annable, and S. N. Young, “Tryptophan, Serotonin and Human Social Behavior,” in Developments in Tryptophan and Serotonin Metabolism, G. Allegri, C. V. L. Costa, E. Ragazzi, H. Steinhart, and L. Varesio, Eds. Boston, MA: Springer US, 2003, pp. 215–224. doi: 10.1007/978-1-4615-0135-0_25.

[39] M. A. Arbib and J.-M. Fellous, “Emotions: from brain to robot,” Trends in Cognitive Sciences, vol. 8, no. 12, pp. 554–561, Dec. 2004, doi: 10.1016/j.tics.2004.10.004.

[40] K. L. Gobrogge, X. Jia, Y. Liu, and Z. Wang, “Neurochemical Mediation of Affiliation and Aggression Associated With Pair-Bonding,” Biol Psychiatry, vol. 81, no. 3, pp. 231–242, Feb. 2017, doi: 10.1016/j.biopsych.2016.02.013.

[41] A. Adell, “Revisiting the role of raphe and serotonin in neuropsychiatric disorders,” J Gen Physiol, vol. 145, no. 4, pp. 257–259, Apr. 2015, doi: 10.1085/jgp.201511389.

[42] R. L. I. Pillai et al., “Examining raphe-amygdala structural connectivity as a biological predictor of SSRI response,” J Affect Disord, vol. 256, pp. 8–16, Sep. 2019, doi: 10.1016/j.jad.2019.05.055.

[43] C. A. Lowry et al., “Serotonergic Systems, Anxiety, and Affective Disorder,” Annals of the New York Academy of Sciences, vol. 1148, no. 1, pp. 86–94, 2008, doi: https://doi.org/10.1196/annals.1410.004.

[44] P. Dayan and Q. J. M. Huys, “Serotonin in Affective Control,” Annu. Rev. Neurosci., vol. 32, no. 1, pp. 95–126, Jun. 2009, doi: 10.1146/annurev.neuro.051508.135607.

[45] Y. Ohmura, T. Izumi, T. Yamaguchi, I. Tsutsui-Kimura, T. Yoshida, and M. Yoshioka, “The Serotonergic Projection from the Median Raphe Nucleus to the Ventral Hippocampus is Involved in the Retrieval of Fear Memory Through the Corticotropin-Releasing Factor Type 2 Receptor,” Neuropsychopharmacology, vol. 35, no. 6, Art. no. 6, May 2010, doi: 10.1038/npp.2009.229.

[46] A. Anand, S. E. Jones, M. Lowe, H. Karne, and P. Koirala, “Resting State Functional Connectivity of Dorsal Raphe Nucleus and Ventral Tegmental Area in Medication-Free Young Adults With Major Depression,” Front Psychiatry, vol. 9, p. 765, 2018, doi: 10.3389/fpsyt.2018.00765.

[47] J. Brakowski et al., “Resting state brain network function in major depression - Depression symptomatology, antidepressant treatment effects, future research,” J Psychiatr Res, vol. 92, pp. 147–159, Sep. 2017, doi: 10.1016/j.jpsychires.2017.04.007.

[48] J. J. Weinstein et al., “Effects of acute tryptophan depletion on raphé functional connectivity in depression,” Psychiatry Res, vol. 234, no. 2, pp. 164–171, Nov. 2015, doi: 10.1016/j.pscychresns.2015.08.015.

[49] J.-P. Hornung, “The human raphe nuclei and the serotonergic system,” Journal of Chemical Neuroanatomy, vol. 26, no. 4, pp. 331–343, Dec. 2003, doi: 10.1016/j.jchemneu.2003.10.002.

[50] L. Schmaal et al., “Subcortical brain alterations in major depressive disorder: findings from the ENIGMA Major Depressive Disorder working group,” Mol Psychiatry, vol. 21, no. 6, pp. 806–812, Jun. 2016, doi: 10.1038/mp.2015.69.

[51] A. Meneses and G. Liy-Salmeron, “Serotonin and emotion, learning and memory,” Reviews in the Neurosciences, vol. 23, no. 5–6, pp. 543–553, Nov. 2012, doi: 10.1515/revneuro-2012-0060.

[52] P. W. Gold, “The organization of the stress system and its dysregulation in depressive illness,” Molecular Psychiatry, vol. 20, no. 1, Art. no. 1, Feb. 2015, doi: 10.1038/mp.2014.163.

[53] E. Baker and F. Lui, “Neuroanatomy, Vagal Nerve Nuclei (Nucleus Vagus),” in StatPearls, Treasure Island (FL): StatPearls Publishing, 2020. Accessed: May 12, 2020. [Online]. Available: http://www.ncbi.nlm.nih.gov/books/NBK545209/

[54] D. A. Groves and V. J. Brown, “Vagal nerve stimulation: a review of its applications and potential mechanisms that mediate its clinical effects,” Neuroscience & Biobehavioral Reviews, vol. 29, no. 3, pp. 493–500, May 2005, doi: 10.1016/j.neubiorev.2005.01.004.

[55] S. J. Sara, “The locus coeruleus and noradrenergic modulation of cognition,” Nat Rev Neurosci, vol. 10, no. 3, pp. 211–223, Mar. 2009, doi: 10.1038/nrn2573.

[56] G. Aston-Jones and B. Waterhouse, “Locus Coeruleus: From Global Projection System to Adaptive Regulation of Behavior,” Brain Res, vol. 1645, pp. 75–78, Aug. 2016, doi: 10.1016/j.brainres.2016.03.001.

[57] S. Bouret and S. J. Sara, “Network reset: a simplified overarching theory of locus coeruleus noradrenaline function,” Trends Neurosci, vol. 28, no. 11, pp. 574–582, Nov. 2005, doi: 10.1016/j.tins.2005.09.002.

[58] G. Aston-Jones and J. D. Cohen, “An integrative theory of locus coeruleus-norepinephrine function: adaptive gain and optimal performance,” Annu Rev Neurosci, vol. 28, pp. 403–450, 2005, doi: 10.1146/annurev.neuro.28.061604.135709.

[59] C. L. Williams, D. Men, and E. C. Clayton, “The effects of noradrenergic activation of the nucleus tractus solitarius on memory and in potentiating norepinephrine release in the amygdala,” Behav Neurosci, vol. 114, no. 6, pp. 1131–1144, Dec. 2000, doi: 10.1037//0735-7044.114.6.1131.

[60] J. T. Coull, C. Büchel, K. J. Friston, and C. D. Frith, “Noradrenergically mediated plasticity in a human attentional neuronal network,” Neuroimage, vol. 10, no. 6, pp. 705–715, Dec. 1999, doi: 10.1006/nimg.1999.0513.

[61] V. Sterpenich et al., “The Locus Ceruleus Is Involved in the Successful Retrieval of Emotional Memories in Humans,” J. Neurosci., vol. 26, no. 28, pp. 7416–7423, Jul. 2006, doi: 10.1523/JNEUROSCI.1001-06.2006.

[62] D. J. H. A. Henssen et al., “Visualizing the trigeminovagal complex in the human medulla by combining ex-vivo ultra-high resolution structural MRI and polarized light imaging microscopy,” Sci Rep, vol. 9, no. 1, p. 11305, Dec. 2019, doi: 10.1038/s41598-019-47855-5.

[63] E. C. Clayton and C. L. Williams, “Adrenergic activation of the nucleus tractus solitarius potentiates amygdala norepinephrine release and enhances retention performance in emotionally arousing and spatial memory tasks,” Behav Brain Res, vol. 112, no. 1–2, pp. 151–158, Jul. 2000, doi: 10.1016/s0166-4328(00)00178-9.

[64] K. Singh et al., “Probabilistic atlas of the lateral parabrachial nucleus, medial parabrachial nucleus, vestibular nuclei complex and medullary viscero-sensory-motor nuclei complex in living humans from 7 Tesla MRI,” bioRxiv, p. 814228, Jan. 2019, doi: 10.1101/814228.

[65] D. A. Groves, E. M. Bowman, and V. J. Brown, “Recordings from the rat locus coeruleus during acute vagal nerve stimulation in the anaesthetised rat,” Neurosci. Lett., vol. 379, no. 3, pp. 174–179, May 2005, doi: 10.1016/j.neulet.2004.12.055.

[66] C. Chen, M. Q. Chen, H. X. Wang, and Y. Chen, “The role of the nucleus tractus solitarius and the locus coeruleus in the abdominal vagal pressor response,” Proc Chin Acad Med Sci Peking Union Med Coll, vol. 4, no. 3, pp. 142–146, 1989.

[67] D. L. Hassert, T. Miyashita, and C. L. Williams, “The Effects of Peripheral Vagal Nerve Stimulation at a Memory-Modulating Intensity on Norepinephrine Output in the Basolateral Amygdala,” Behavioral Neuroscience, vol. 118, no. 1, pp. 79–88, 2004, doi: 10.1037/0735-7044.118.1.79.

[68] P. Rutecki, “Anatomical, physiological, and theoretical basis for the antiepileptic effect of vagus nerve stimulation,” Epilepsia, vol. 31 Suppl 2, pp. S1-6, 1990, doi: 10.1111/j.1528-1157.1990.tb05843.x.

[69] T. R. Henry, R. A. E. Bakay, P. B. Pennell, C. M. Epstein, and J. R. Votaw, “Brain blood-flow alterations induced by therapeutic vagus nerve stimulation in partial epilepsy: II. prolonged effects at high and low levels of stimulation,” Epilepsia, vol. 45, no. 9, pp. 1064–1070, Sep. 2004, doi: 10.1111/j.0013-9580.2004.03104.x.

[70] A. Barnes, R. Duncan, J. A. Chisholm, K. Lindsay, J. Patterson, and D. Wyper, “Investigation into the mechanisms of vagus nerve stimulation for the treatment of intractable epilepsy, using 99mTc-HMPAO SPET brain images,” Eur J Nucl Med Mol Imaging, vol. 30, no. 2, pp. 301–305, Feb. 2003, doi: 10.1007/s00259-002-1026-8.

[71] R. Ruffoli, F. S. Giorgi, C. Pizzanelli, L. Murri, A. Paparelli, and F. Fornai, “The chemical neuroanatomy of vagus nerve stimulation,” J. Chem. Neuroanat., vol. 42, no. 4, pp. 288–296, Dec. 2011, doi: 10.1016/j.jchemneu.2010.12.002.

[72] F. Fornai, R. Ruffoli, F. S. Giorgi, and A. Paparelli, “The role of locus coeruleus in the antiepileptic activity induced by vagus nerve stimulation,” European Journal of Neuroscience, vol. 33, no. 12, pp. 2169–2178, 2011, doi: 10.1111/j.1460-9568.2011.07707.x.

[73] K. T. Beier et al., “Circuit Architecture of VTA Dopamine Neurons Revealed by Systematic Input-Output Mapping,” Cell, vol. 162, no. 3, pp. 622–634, Jul. 2015, doi: 10.1016/j.cell.2015.07.015.

[74] Ó. Arias-Carrión and E. Pöppel, “Dopamine, learning, and reward-seeking behavior,” Acta Neurobiologiae Experimentalis, vol. 67, no. 4, pp. 481–488, 2007.

[75] A. Alcaro, R. Huber, and J. Panksepp, “Behavioral functions of the mesolimbic dopaminergic system: An affective neuroethological perspective,” Brain Research Reviews, vol. 56, no. 2, pp. 283–321, Dec. 2007, doi: 10.1016/j.brainresrev.2007.07.014.

[76] M. L. Settell et al., “Functional Circuitry Effect of Ventral Tegmental Area Deep Brain Stimulation: Imaging and Neurochemical Evidence of Mesocortical and Mesolimbic Pathway Modulation,” Front Neurosci, vol. 11, p. 104, 2017, doi: 10.3389/fnins.2017.00104.

[77] S. J. Russo and E. J. Nestler, “The brain reward circuitry in mood disorders,” Nature Reviews Neuroscience, vol. 14, no. 9, Art. no. 9, Sep. 2013, doi: 10.1038/nrn3381.

[78] L. Lu, B. Hope, J. Dempsey, S. Liu, J. Bossert, and Y. Shaham, “Central amygdala ERK signaling pathway is critical to incubation of cocaine craving,” Nature neuroscience, vol. 8, pp. 212–9, Mar. 2005, doi: 10.1038/nn1383.

[79] Chi Yiu Yim and G. J. Mogenson, “Response of nucleus accumbens neurons to amygdala stimulation and its modification by dopamine,” Brain Research, vol. 239, no. 2, pp. 401–415, May 1982, doi: 10.1016/0006-8993(82)90518-2.

[80] J. S. Brog, A. Salyapongse, A. Y. Deutch, and D. S. Zahm, “The patterns of afferent innervation of the core and shell in the ‘Accumbens’ part of the rat ventral striatum: Immunohistochemical detection of retrogradely transported fluoro-gold,” Journal of Comparative Neurology, vol. 338, no. 2, pp. 255–278, 1993, doi: https://doi.org/10.1002/cne.903380209.

[81] P. W. Kalivas, “Neurotransmitter regulation of dopamine neurons in the ventral tegmental area,” Brain Res Brain Res Rev, vol. 18, no. 1, pp. 75–113, Apr. 1993, doi: 10.1016/0165-0173(93)90008-n.

[82] J. A. Hosp et al., “Ventral tegmental area connections to motor and sensory cortical fields in humans,” Brain Struct Funct, vol. 224, no. 8, pp. 2839–2855, Nov. 2019, doi: 10.1007/s00429-019-01939-0.

[83] V. A. Coenen et al., “The anatomy of the human medial forebrain bundle: Ventral tegmental area connections to reward-associated subcortical and frontal lobe regions,” NeuroImage: Clinical, vol. 18, pp. 770–783, Jan. 2018, doi: 10.1016/j.nicl.2018.03.019.

[84] V. A. Coenen et al., “MEDIAL FOREBRAIN BUNDLE STIMULATION AS A PATHOPHYSIOLOGICAL MECHANISM FOR HYPOMANIA IN SUBTHALAMIC NUCLEUS DEEP BRAIN STIMULATION FOR PARKINSON’S DISEASE,” Neurosurgery, vol. 64, no. 6, pp. 1106–1115, Jun. 2009, doi: 10.1227/01.NEU.0000345631.54446.06.

[85] J. M. Anthofer, K. Steib, C. Fellner, M. Lange, A. Brawanski, and J. Schlaier, “DTI-based deterministic fibre tracking of the medial forebrain bundle,” Acta Neurochir, vol. 157, no. 3, pp. 469–477, Mar. 2015, doi: 10.1007/s00701-014-2335-y.

[86] T. R. Wang, S. Moosa, R. F. Dallapiazza, W. J. Elias, and W. J. Lynch, “Deep brain stimulation for the treatment of drug addiction,” Neurosurg Focus, vol. 45, no. 2, p. E11, 2018, doi: 10.3171/2018.5.FOCUS18163.

[87] D. B. Vyas et al., “Deep Brain Stimulation for Chronic Cluster Headache: A Review,” Neuromodulation, vol. 22, no. 4, pp. 388–397, Jun. 2019, doi: 10.1111/ner.12869.

[88] R. R. Rozeske et al., “Prefrontal-Periaqueductal Gray-Projecting Neurons Mediate Context Fear Discrimination,” Neuron, vol. 97, no. 4, pp. 898–910.e6, Feb. 2018, doi: 10.1016/j.neuron.2017.12.044.

[89] C. Silva and N. McNaughton, “Are periaqueductal gray and dorsal raphe the foundation of appetitive and aversive control? A comprehensive review,” Prog Neurobiol, vol. 177, pp. 33–72, Jun. 2019, doi: 10.1016/j.pneurobio.2019.02.001.

[90] M. Ezra, O. K. Faull, S. Jbabdi, and K. T. Pattinson, “Connectivity-based segmentation of the periaqueductal gray matter in human with brainstem optimized diffusion MRI,” Human Brain Mapping, vol. 36, no. 9, pp. 3459–3471, 2015, doi: https://doi.org/10.1002/hbm.22855.

[91] D. T. George, R. Ameli, and G. F. Koob, “Periaqueductal Gray Sheds Light on Dark Areas of Psychopathology,” Trends Neurosci, vol. 42, no. 5, pp. 349–360, May 2019, doi: 10.1016/j.tins.2019.03.004.

[92] S. L. F. Owen et al., “Pre-operative DTI and probabilisitic tractography in four patients with deep brain stimulation for chronic pain,” J Clin Neurosci, vol. 15, no. 7, pp. 801–805, Jul. 2008, doi: 10.1016/j.jocn.2007.06.010.

[93] E. Sillery et al., “Connectivity of the human periventricular—periaqueductal gray region,” Journal of Neurosurgery, vol. 103, no. 6, pp. 1030–1034, Dec. 2005, doi: 10.3171/jns.2005.103.6.1030.

[94] Sims-Williams H et al., “Deep brain stimulation of the periaqueductal gray releases endogenous opioids in humans,” NeuroImage, vol. 146, Feb. 2017, doi: 10.1016/j.neuroimage.2016.08.038.

[95] R. M. Levy, S. Lamb, and J. E. Adams, “Treatment of chronic pain by deep brain stimulation: long term follow-up and review of the literature,” Neurosurgery, vol. 21, no. 6, pp. 885–893, Dec. 1987, doi: 10.1227/00006123-198712000-00017.

[96] R. G. Bittar et al., “Deep brain stimulation for pain relief: a meta-analysis,” J Clin Neurosci, vol. 12, no. 5, pp. 515–519, Jun. 2005, doi: 10.1016/j.jocn.2004.10.005.

[97] A. Bari, T. Niu, J.-P. Langevin, and I. Fried, “Limbic neuromodulation: implications for addiction, posttraumatic stress disorder, and memory,” Neurosurg. Clin. N. Am., vol. 25, no. 1, pp. 137–145, Jan. 2014, doi: 10.1016/j.nec.2013.08.004.

[98] H. Akram et al., “Ventral tegmental area deep brain stimulation for refractory chronic cluster headache,” Neurology, vol. 86, no. 18, pp. 1676–1682, 03 2016, doi: 10.1212/WNL.0000000000002632.

[99] C. Thomas et al., “Anatomical accuracy of brain connections derived from diffusion MRI tractography is inherently limited,” Proc. Natl. Acad. Sci. U.S.A., vol. 111, no. 46, pp. 16574–16579, Nov. 2014, doi: 10.1073/pnas.1405672111.

[100] K. G. Schilling et al., “Limits to anatomical accuracy of diffusion tractography using modern approaches,” Neuroimage, vol. 185, pp. 1–11, Jan. 2019, doi: 10.1016/j.neuroimage.2018.10.029.

[101] M. Kinoshita et al., “Fiber-tracking does not accurately estimate size of fiber bundle in pathological condition: initial neurosurgical experience using neuronavigation and subcortical white matter stimulation,” Neuroimage, vol. 25, no. 2, pp. 424–429, Apr. 2005, doi: 10.1016/j.neuroimage.2004.07.076.

[102] A. Alhourani and R. M. Richardson, “Inherent limitations of tractography for accurate connectivity maps,” Neurosurgery, vol. 76, no. 4, pp. N11-12, Apr. 2015, doi: 10.1227/01.neu.0000462692.36374.1a.

[103] M. A. AbuAlrob and P. Tadi, “Neuroanatomy, Nucleus Solitarius,” in StatPearls, Treasure Island (FL): StatPearls Publishing, 2020. Accessed: Oct. 09, 2020. [Online]. Available: http://www.ncbi.nlm.nih.gov/books/NBK549831/

[104] L. D. Hachem, S. M. Wong, and G. M. Ibrahim, “The vagus afferent network: emerging role in translational connectomics,” Neurosurg Focus, vol. 45, no. 3, p. E2, 2018, doi: 10.3171/2018.6.FOCUS18216.

[105] E. E. Benarroch, “The central autonomic network: functional organization, dysfunction, and perspective,” Mayo Clinic Proceedings, vol. 68, no. 10, pp. 988– 1001, Oct. 1993, doi: 10.1016/s0025-6196(12)62272-1.

[106] O. Lindvall, A. Björklund, A. Nobin, and U. Stenevi, “The adrenergic innervation of the rat thalamus as revealed by the glyoxylic acid fluorescence method,” Journal of Comparative Neurology, vol. 154, no. 3, pp. 317–347, 1974, doi: 10.1002/cne.901540307.

[107] E. E. Benarroch, “Locus coeruleus,” Cell Tissue Res., vol. 373, no. 1, pp. 221– 232, Jul. 2018, doi: 10.1007/s00441-017-2649-1.

[108] F. Hernández-Vázquez, J. Garduño, and S. Hernández-López, “GABAergic modulation of serotonergic neurons in the dorsal raphe nucleus,” Rev Neurosci, vol. 30, no. 3, pp. 289–303, 24 2019, doi: 10.1515/revneuro-2018-0014.

[109] R. W. Roosevelt, D. C. Smith, R. W. Clough, R. A. Jensen, and R. A. Browning, “Increased extracellular concentrations of norepinephrine in cortex and hippocampus following vagus nerve stimulation in the rat,” Brain Res., vol. 1119, no. 1, pp. 124–132, Nov. 2006, doi: 10.1016/j.brainres.2006.08.048.

[110] R. Browning, K. Clark, D. Naritoku, D. Smith, and R. Jensen, “Loss of anticonvulsant effect of vagus nerve stimulation in the pentylenetetrazol seizure model following treatment with 6-hydroxydopamine or 5,7-dihydroxytryptamine. Browning RA, Clark KB, Naritoku DK, Smith DC, Jensen RA:,” Soc Neurosci Abstr, vol. 23, no. 2424, 1997.

